# DnaJB1 chaperone inhibits tau aggregation by recognizing its N-terminus

**DOI:** 10.1101/2025.10.02.680111

**Authors:** Pawel M. Wydorski, Paulina Macierzynska, Sofia Bali, Jaime Vaquer-Alicea, Lukasz A. Joachimiak

## Abstract

A network of protein folding and degradation machineries maintains protein homeostasis by preventing the accumulation of misfolded proteins and by facilitating their clearance. These systems are also crucial for the inhibition of protein aggregation in neurodegenerative diseases where misfolded proteins often aggregate into β-rich amyloid fibrils. How these machineries selectively recognize pathological aggregates over normal conformations remains unclear. Here, we present the molecular logic for how a Hsp70 co-chaperone from the J-domain protein family, DnaJB1, binds pathological aggregates of the microtubule-associated protein tau through the recognition of the flexible N-terminus that comprises the disordered fuzzy coat of fibrils. We show that this interaction contributes to the regulation of tau assembly in cellular models of tau aggregation and depends on the presence of the negatively charged residues. We determined that DnaJB1 inhibits tau aggregation in vitro through these interactions, and found that this weak, transient binding can be enhanced by the presence of polyanionic factors such as heparin. As prospective client-binding sites, we identified the charged hinge between the two β-sandwich C-terminal domains I and II, as well as the conserved J-domain of this chaperone. This work presents novel biochemical and structural insights into how the molecular chaperone DnaJB1 recognizes full-length forms of tau protein in a pathological context.

## Introduction

The microtubule-associated protein tau accumulates as β-sheet-rich amyloid fibrils in a group of neurodegenerative diseases, including Alzheimer’s disease (Goedert et al., 1988; Wischik et al., 1988; V. M.-Y. Lee et al., 1991). The precise trigger of tau fibrilization in human brains is still unknown. Tau is a highly soluble protein in monomeric form and doesn’t aggregate easily in the context of in vitro studies without utilization of mutations and/or additional factors (Barghorn & Mandelkow, 2002). Mutations in tau lead to frontotemporal dementia (FTD), but studies utilizing these pathology-associated mutations have shown that these mutations alone often do not lead to spontaneous tau aggregation. To date, the only mutant known to drive spontaneous tau aggregation in vitro and in cells is the substitution from serine to phenylalanine at position 320 (Strang et al., 2018; D. Chen et al., 2023). Alternative ways to promote tau assembly into filaments combine incubation of full-length (FL) tau or its fragments at high concentration in buffers with defined ionic strengths (Lövestam et al., 2022, 2025). These new approaches to control tau’s aggregation into amyloids overcome the reliance on incubating tau with polyanionic inducers such as heparin, RNA or arachidonic acid (Pérez et al., 1996; Montgomery et al., 2023). Initiation of tau fibrilization with heparin shortens the aggregation lag time by reducing the cationic repulsion between the positively charged repeat domains of tau monomer (Zhao et al., 2017) and increasing the accessibility of the amyloid motifs localized in repeat domains 2 and 3 of 4R tau (Friedhoff et al., 1998; Von Bergen et al., 2000, 2001; D. Chen et al., 2019). Initial studies suggested a physiological role of heparan sulphate in tau fibrilization (Goedert et al., 1996). However, recent cryo-EM structures uncovered polymorphism of heparin-induced aggregates that bear little structural similarity to ex vivo fibrils from any tauopathy (Zhang et al., 2019). This confounds the physiological relevance of studies that used heparin-induced tau fibrils to identify cellular factors, like molecular chaperones, that control different steps of tau assembly/disassembly, or to discover drug-like molecules that could slow amyloid formation. More mechanistic studies are needed to better understand the suitability of using heparin as the inducer of tau fibrilization.

The Hsp70/J-domain protein (JDP) system plays an important role in ensuring correct protein folding in cells (Rosenzweig et al., 2019). In humans, JDP family contains nearly 50 members with diverse activities that range from regulation of protein folding/refolding, complex assembly, protein import into organelles, to stress response (Kampinga & Craig, 2010). Importantly, JDPs directly bind to substrates and recruit them to the Hsp70 system for ATP-dependent processing, thus imparting specificity to the generic binding capability of Hsp70s. Members of this protein family across its three classes (A, B, and C) have been found to be the modulators of aggregation for many proteins associated with neurodegenerative diseases, such as α-synuclein (aSyn), huntingtin, tau, and TDP-43 (Pemberton et al., 2011; Wentink et al., 2020; Muchowski et al., 2000; Ayala Mariscal et al., 2022; Irwin et al., 2021; Hou et al., 2021; Abayev-Avraham et al., 2023). In the context of tau, DnaJA1 and DnaJA2 were shown to bind monomeric species of tau with DnaJA2 also binding heparin-induced tau fibrils composed of the repeat domains of tau (tauRD) (Mok et al., 2018; Irwin et al., 2021). In C class, DnaJC7 preferentially binds to the native WT FL tau monomer and modulates seeding in cell models of tau aggregation (Hou et al., 2021; Perez et al., 2023), and DnaJC5 chaperone has been reported to increase extracellular tau release (Fontaine et al., 2016; Xu et al., 2018).

Among class B JDPs, studies report that DnaJB1 recognizes mutant and heparin-incubated tauRD monomers, but not native wild-type (WT) FL tau (Irwin et al., 2021). Additionally, DnaJB1, as well as DnaJB4 can bind to heparin-induced fibrils composed of tauRD and FL tau (Nachman et al., 2020; Irwin et al., 2021). DnaJB1 has been shown to interact in vitro with multiple other aggregation-prone proteins including huntingtin and TDP-43 (Ayala Mariscal et al., 2022; Carrasco et al., 2023). However, the largest body of studies was conducted in the context of recognition—and most notably—disassembly of aSyn fibrils, a protein associated with pathologies such as multiple system atrophy and Parkinson’s disease. It was found that DnaJB1 can recognize and bind to the acidic C-terminal region of aSyn monomers which remains unstructured in the fibril form. This leads to the recruitment of Hsp70 and Hsp110 to the fibril with subsequent disaggregation via entropic pulling (Gao et al., 2015; Wentink et al., 2020; Monistrol et al., 2025).

The role of the fuzzy coat in tau fibrils became of interest, for example, with recent results postulating its role in determining tau’s seeding capacity (Kasen et al., 2025). This underlies the necessity of consideration of the full-length protein in studying tau fibrilization, and how other cellular factors may interact with these aggregates. To date, the biochemical or structural information on how molecular chaperones recognize the full-length forms of tau protein remains limited. In this work, we show how the DnaJB1 chaperone binds to N-terminal motifs allowing it to act on tau monomer as well as pathological tau fibrils. The residues composing these motifs are predominantly negatively charged and hydrophobic, and truncation of tau from the N-terminus at these regions reveals changes in the dependence of DnaJB1 on tau assembly in cellular models of tau aggregation. In vitro heparin-less aggregation studies using FL tau and tauRD show that DnaJB1 efficiently suppresses aggregation of FL tau but not tauRD. Paradoxically, parallel experiments in the presence of heparin show that DnaJB1 does not prevent FL tau or tauRD aggregation. Using binding assays, we demonstrate that DnaJB1’s recruitment to tau can be enhanced by the presence of the polyanions such as heparin. In addition to the previously reported client-binding sites localized on the DnaJB1’s C-terminal domains I and II (CTDI and CTDII), we report new interactions that depend on the chaperone’s J-domain (JD). JD has been previously shown to be responsible for the recruitment of the chaperone-client complex to the Hsp70 machinery for further processing, as well as for stimulating Hsp70’s ATPase activity. Our findings yield new insights into the molecular recognition between DnaJB1-tau and uncover potential shortfalls of heparin utilization as a fibrilization inducer, as we identified competitive binding modes that could occlude physiologically relevant molecular interactions. These results help to deconvolute previously unknown mechanisms of interactions between members of the protein homeostasis network and aggregation-prone proteins.

## Results

### Truncation of tau at the N-terminus alters its aggregation dependence on DnaJB1

In the context of aSyn fibrils, DnaJB1 was found to bind an acidic region at the C-terminus of the protein which remains disordered in aSyn fibrils (Wentink et al., 2020). Studies on tau showed that DnaJB1 and Hsp70 machinery can disassemble recombinant tau fibrils composed of various tau isoforms (Nachman et al., 2020). Based on these results, we hypothesized that DnaJB1 recognizes disordered elements in the tau sequence present in the fuzzy coat of tau fibrils (Fitzpatrick et al., 2017). In tau fibrils, the N-terminal region of tau (N-terminal to the repeat domains) is not incorporated in any known core of tau fibrils and is predominantly composed of negatively charged residues (Figure 1A; with net charge -7 at physiological pH for residues 1-243). First, we tested the role of DnaJB1 in tau fibrilization utilizing a cellular model of tau aggregation, taking advantage of a photoconvertible fluorophore that enables FRET-based quantification of tau aggregate formation (D. Chen et al., 2023). Introduction of exogenous aggregates into tau biosensor cells using lipofectamine induces aggregation of the intracellular tau which can be quantified using FRET (Holmes et al., 2014; Furman et al., 2015). First, we created two different DnaJB1 expression contexts in HEK293T cells: (1) DnaJB1-mCerulean3 overexpression (OE) and (2)

**Figure 1.**
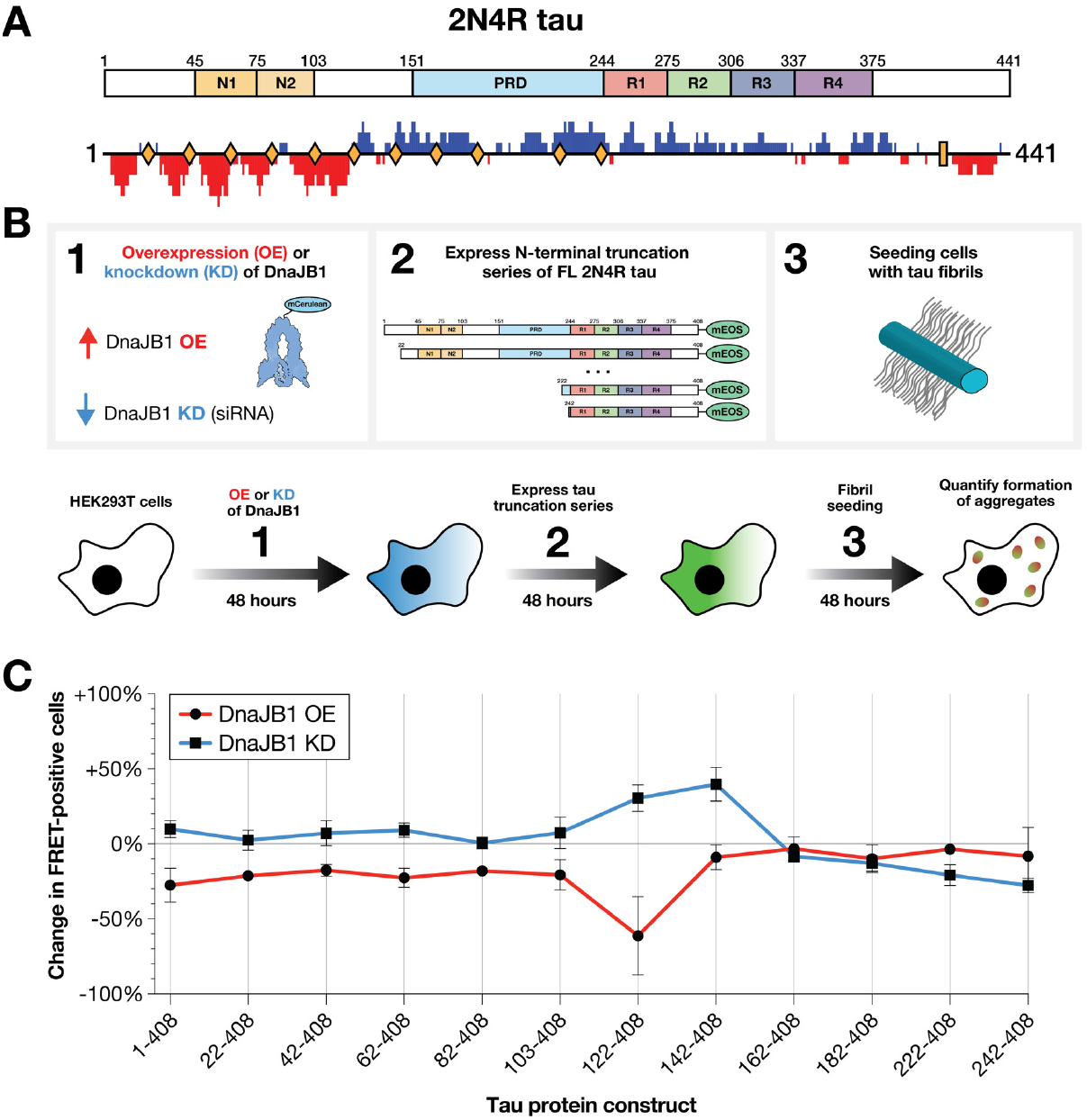
DnaJB1 recognizes the acidic region of the N-terminus of microtubule-associated protein tau. **(A)** The domain architecture of 2N4R isoform of the microtubule-associated protein tau. N1 and N2 are two N-terminal domains, PRD is Proline-Rich Domain, and R1-4 are Repeat Domains from 1 to 4. Below is the calculated linear net charge per residue (NCPR) using CIDER (Holehouse et al., 2017) with blob size 10. Yellow diamonds represent ranges of the N-terminal truncations of 2N4R tau used in this study. **(B)** Schematic of the experimental setup using cellular model of tau aggregation. HEK293T cells were first transfected with either DnaJB1-mCerulean3 plasmid or siRNA against human DnaJB1, 24 hours after plating. After 48 hours, cells were further transected with plasmids containing different N-terminal truncations of 2N4R isoform of tau protein with C-terminal mEOS3.2 fluorescent tag. After 48 hours of expression, the cells were seeded with recombinant tau fibrils and tau aggregation was measured as a FRET signal via flow cytometry 48 hours after seeding with recombinant tau fibrils. **(C)** Percentage change in tau aggregation (measured as a FRET signal from photoconverted mEOS3.2 fluorescent tag) of different N-terminal truncations of tau protein in cell biosensors. Red line corresponds to DnaJB1 overexpression context and blue line corresponds to the DnaJB1 knockdown. Data are shown as averages with error bars representing SEM for 3 biological replicates.

DnaJB1 knockdown (KD) using siRNA (Fig. 1B, Supplementary Fig. 1AB). A series of N-terminal truncations of tau in 20 amino acid increments were generated as C-terminal fusions to a photoconvertible and FRET-compatible mEOS3.2 fluorescent tag. This panel of tau variants was co-expressed with DnaJB1-mCerulean3 (i.e. DnaJB1 OE) or with siRNA targeting DnaJB1 (i.e. DnaJB1 KD). These cells were then seeded with exogenous tau aggregates to initiate aggregation of each tau-mEOS3.2 variant. Seeding activity in each condition was quantified by FRET using flow cytometry, with observed changes referencing to mCerulean3-only overexpression or scramble sequence siRNA controls for OE and KD, respectively (Fig. 1C, Supplementary Fig. 1CD, see also Materials and methods). OE of DnaJB1 inhibited tau aggregate formation in longer constructs that maintained the majority of the acidic 120 N-terminal amino acids of 2N4R isoform of tau, but OE of DnaJB1 on shorter tau constructs had no effect (Fig. 1C). Noticeably, for the six longest among the tested constructs, removing 20 residues up to residue 103 resulted in up to a 30% reduction in tau seeding. Removing the next 20 residues (103-122) had a large effect on tau aggregation in DnaJB1 OE context, whereas for the shortest tested tau constructs (lacking first 142 residues and shorter), chaperone overexpression no longer resulted in reduction of aggregation. To complement the OE study, we designed parallel experiments to test reduction in DnaJB1 and used the siRNA approach to knockdown expression of this chaperone. Optimization of siRNA conditions targeting DnaJB1 yielded 65% reduction in protein levels (Supplementary Fig. 1B). KD of DnaJB1 using siRNA only modestly increased tau seeding, and notably, the effects mirror those observed with DnaJB1 OE where KD increased seeding for N-terminal truncations with the largest increase for the 142-408 construct (Fig. 1C). The most pronounced change in seeding (both OE and KD) is observed with truncation variants of tau that shift the charge distribution of 2N4R tau from predominantly negative to positive between residues 120-140 (Fig. 1AC), highlighting the role of the negatively charged N-terminus in enabling DnaJB1’s recruitment to the fibril, similarly to what has been reported for aSyn (Wentink et al., 2020; Jäger et al., 2024).

### Inhibition of tau aggregation by DnaJB1 is dependent on the presence of fibril’s fuzzy coat

The effect of DnaJB1 on tau aggregation kinetics in vitro have been previously reported but required the co-aggregation of tau and heparin with the chaperone (Sahara et al., 2007; Irwin et al., 2021). A recent study reported that tau encoding glutamic acid phosphomimetic substitutions was shown to form fibrils in the absence of polyanions but rather utilizing potassium citrate (Lövestam et al., 2025). Using these conditions, we produced in vitro fibrils using a tau repeat domain (tauRD, residues 244-380 in 2N4R isoform) construct that is missing the N-terminal fuzzy coat and FL 2N4R tau encoding a S320F mutation that we have previously shown to aggregate spontaneously, while maintaining the length of the disordered fuzzy coat of the fibril (Strang et al., 2018; D. Chen et al., 2023). Taking advantage of these new systems, we tested DnaJB1’s impact on tau aggregation in heparin-less conditions for FL tau and tauRD (Fig. 2A). We monitored aggregation kinetics using a thioflavin T (ThT) fluorescence aggregation assay while titrating DnaJB1 (Fig. 2B, Supplementary Fig. 2A). We found concentration-dependent inhibition of FL tau fibrilization by DnaJB1, with 5:1 molar ratio of tau to chaperone being sufficient to nearly abolish ThT-positive fibril formation (Fig. 2BC). Fibrilization kinetics of tauRD in heparin-less conditions were significantly delayed by DnaJB1 but the decrease of ThT fluorescence signal at the endpoints was more variable (Fig. 2BC). Seeding of the FL tau reactions co-incubated with DnaJB1 correlated well with the in vitro ThT fluorescence aggregation assay, and at higher concentrations of the chaperone (starting at 5:1) seeding activity dropped to lipofectamine control levels. Interestingly, tauRD co-incubated with DnaJB1 led to increases in the percentage of cells with FRET-positive tau inclusions in a chaperone concentration-dependent manner (Fig. 2DE, Supplementary Fig. 2C). This observation is in line with the previous findings that DnaJB1 in concert with Hsp70 machinery can lead to the development of more seeding-competent species of tau and aSyn fibrils (Nachman et al., 2020; Tittelmeier et al., 2022). Our results demonstrate that the N-terminus of tau is required for DnaJB1 to inhibit tau fibrilization and seed formation as seen for FL tau aggregation in heparin-less conditions. By contrast, heparin-less aggregation of tauRD in the presence of DnaJB1, delays kinetics as a function of chaperone concentration but produce more active species in cell-based seeding assays suggesting the emergence of seeds that are not ThT positive but retain seeding activity (Fig. 2DE, Supplementary Fig. 2C).

**Figure 2.**
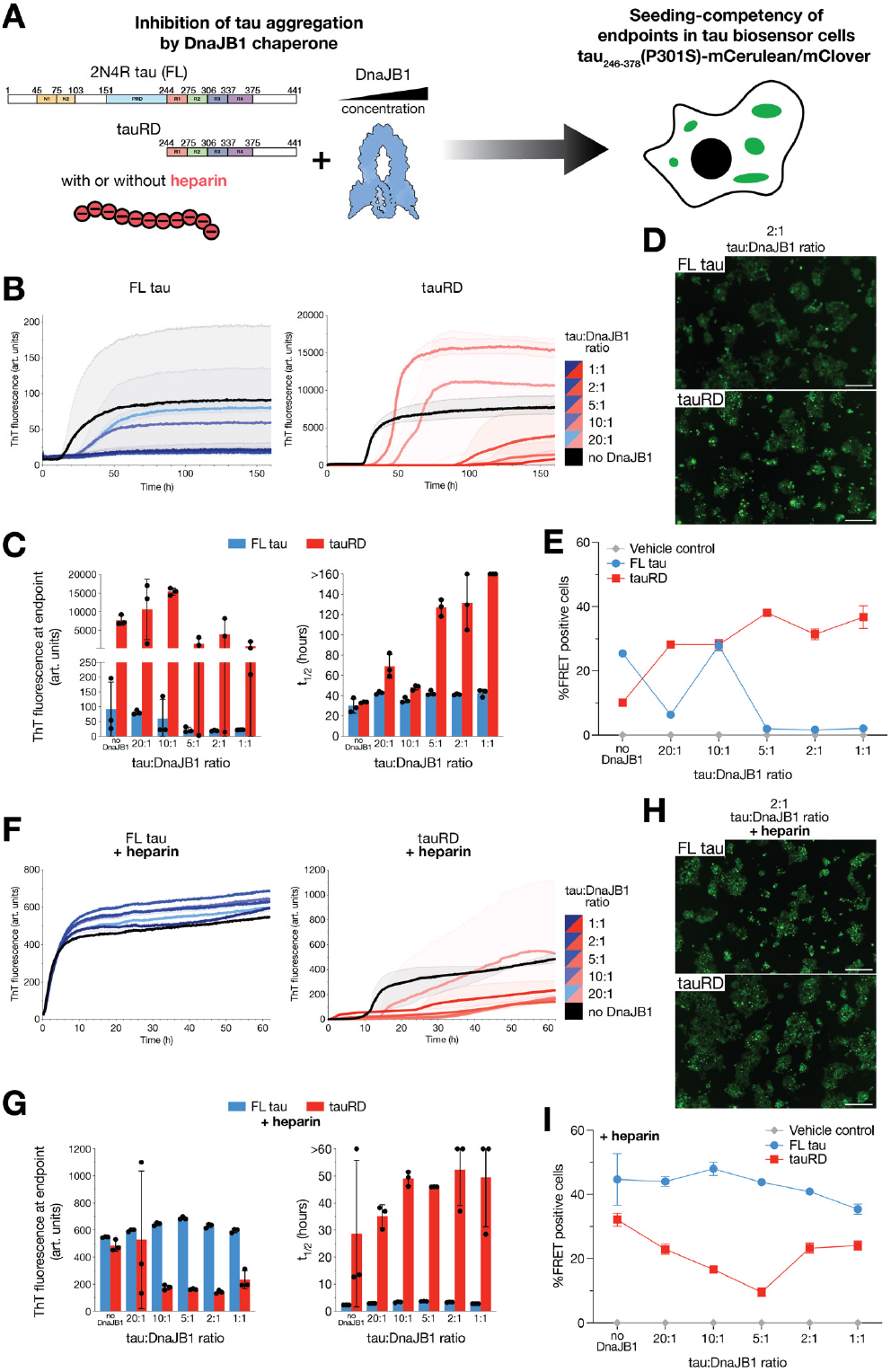
Aggregation of tau in presence of DnaJB1 leads to less seeding-competent species and DnaJB1’s impact is dependent on tau’s N-terminus. **(A)** Schematic of the experimental design. Full-length (FL) or repeat domain (tauRD) constructs of tau protein were aggregated with different concentrations of DnaJB1 chaperone in heparin-less or heparin-containing conditions. Endpoints of aggregation assay were then added to cell biosensors of tau aggregation. **(B)** Thioflavin T (ThT) fluorescence signals of FL S320F tau (left, blue) and WT tauRD (right, red) incubated in absence of DnaJB1 (black) or with different concentrations of DnaJB1 (color gradients) in 100 mM potassium phosphate, 400 mM potassium citrate, 4 mM TCEP, pH 7.2. Each condition with different chaperone concentration was performed in triplicate. The data are shown as solid and transparent lines that represent the average and range across replicates, respectively. **(C)** Parameters from ThT fluorescence aggregation assay shown at (B) presented as the fluorescence signal at the endpoint (left) and t_1/2_ max fit to each aggregation curve (right) for FL S320F tau (blue) and WT tauRD (red). Data are plotted as an average with a SD across three replicates. **(D)** Representative fluorescent microscope images of cell seeding experiments of ThT aggregation assay endpoints from (B) for 2:1 of tau:DnaJB1 ratio condition for aggregation of FL S320F tau (top) and tauRD (bottom) in heparin-less conditions. tauRD biosensors were seeded with 100 nM final concentration of the ThT assay endpoints and after 48 hours the development of tau inclusions was investigated. Scale bars represent 200 μm. **(E)** Quantification of tau inclusion formation when tau biosensor cells were seeded with 100 nM of the ThT assay endpoints of FL S320F tau (blue) or WT tauRD (red) incubated with different concentrations of DnaJB1 in heparin-less buffer conditions. Data are plotted as an average with a SD across three independent transfections. **(F)** ThT fluorescence signals of FL S320F tau (left, blue) and WT tauRD (right, red) incubated in absence of DnaJB1 (black) or with different concentrations of DnaJB1 (color gradients) in 1X PBS, 10 mM DTT, pH 7.4 in presence of equimolar concentration of heparin. Each condition with different chaperone concentration was performed in triplicate. The data are shown as solid and transparent lines that represent the average and range across replicates, respectively. **(G)** Parameters from ThT fluorescence aggregation assay shown at (F) presented as the fluorescence signal at the endpoint (left) and t_1/2_ max fit to each aggregation curve (right) for FL S320F tau (blue) and WT tauRD (red). Data are plotted as an average with SD across three replicates. **(H)** Representative fluorescent microscope images of cell seeding experiments of ThT aggregation assay endpoints from (F) for 2:1 of tau:DnaJB1 ratio condition for aggregation of FL S320F tau (top) and tauRD (bottom) in heparin present conditions. Tau biosensors were seeded with 100 nM final concentration of the ThT assay endpoints and after 48 hours the development of tau inclusions was investigated. Scale bars represent 200 μm. **(I)** Quantification of tau inclusion formation when tau biosensor cells were seeded with 100 nM of the ThT assay endpoints of FL S320F tau (blue) or WT tauRD (red) incubated with different concentrations of DnaJB1 in heparin present conditions. Data are plotted as an average with SD across three independent transfections.

When tau and heparin were co-incubated with DnaJB1 in phosphate buffer in the presence of heparin, we observed a similar dose-dependent effect of this chaperone on aggregation of tauRD, but we did not meansure a reduction in aggregation of FL tau (Fig. 2F). Surprisingly, co-incubation of FL tau and heparin with DnaJB1 led to an increase in ThT fluorescence signal when the chaperone was present (Fig. 2FG). These results may be explained by DnaJB1 itself contributing to the ThT-positive fluorescence signal, consistent with the emergence of β-sheet-rich structures, with heparin incubation leading to stronger effects (Supplementary Fig. 2D). However, these DnaJB1 species did not induce tau aggregation in our cellular model (Supplementary Fig. 2CE). When we tested the endpoints of the in vitro reactions on tau aggregation biosensors, we found that heparin was more effective in inducing formation of the seeding-competent tau species than potassium citrate. However, the effect of the DnaJB1 chaperone in lowering the emergence of tau seeds is lost for both FL tau and tauRD when incubated with heparin (Fig. 2HI, Supplementary Fig. 2C).

### Heparin enhances DnaJB1’s recruitment to tau monomers

Motivated by the effects of DnaJB1 on inhibiting FL tau aggregation, we quantified the interactions between DnaJB1 and tau. We decided to measure the binding affinities of this chaperone to different tau constructs using Isothermal Calorimetry (ITC) (Fig. 3A). Surprisingly, we did not detect changes in heat released when titrating FL tau with DnaJB1 in the tested concentration range (up to 200 μM) (Fig. 3B). Next, we tested whether DnaJB1 binding may depend on changes in local conformational changes in FL tau monomer that arise from different disease-associated point mutations as shown previously (Hou et al., 2021; Irwin et al., 2021). Using Microscale Thermophoresis (MST) with fluorescently labeled DnaJB1, we found no substantial differences in binding of various pathology-associated point mutants of 2N4R tau with mutations localized inside or outside of the repeat domain, with only weak, likely non-specific, binding detected that did not allow for affinity estimation (Supplementary Fig. 3BC). No differences in binding efficiencies were found between WT and S320F FL tau monomers, suggesting that the tau aggregation inhibition observed in previous experiments is not caused by the increased DnaJB1 binding to S320F FL tau. To address the possibility that the intra-molecular interactions of tau monomer, known as the ‘paperclip’ conformation of tau in solution, decrease the accessibility of the binding sites for the chaperone, we next tested whether DnaJB1 can interact with the isolated N-terminal fragment of tau (here referred to as tau_1-243_) or the tau repeat domain (tauRD), and similarly to FL, we were not able to detect binding in the concentration range tested (Fig. 3B). Prior studies suggested that DnaJB1 binding to aSyn strongly relies on ionic strength (Monistrol et al., 2025). To test this, and whether interactions between DnaJB1 and tau_1-243_ depend on electrostatics, we also measured DnaJB1:tau_1-243_ binding in buffers with different ionic strengths but found no increase in binding (Supplementary Fig. 3A). We conclude that tau’s disease-associated point mutations, its truncations, or changes in buffer composition do not enhance DnaJB1 binding in vitro.

**Figure 3.**
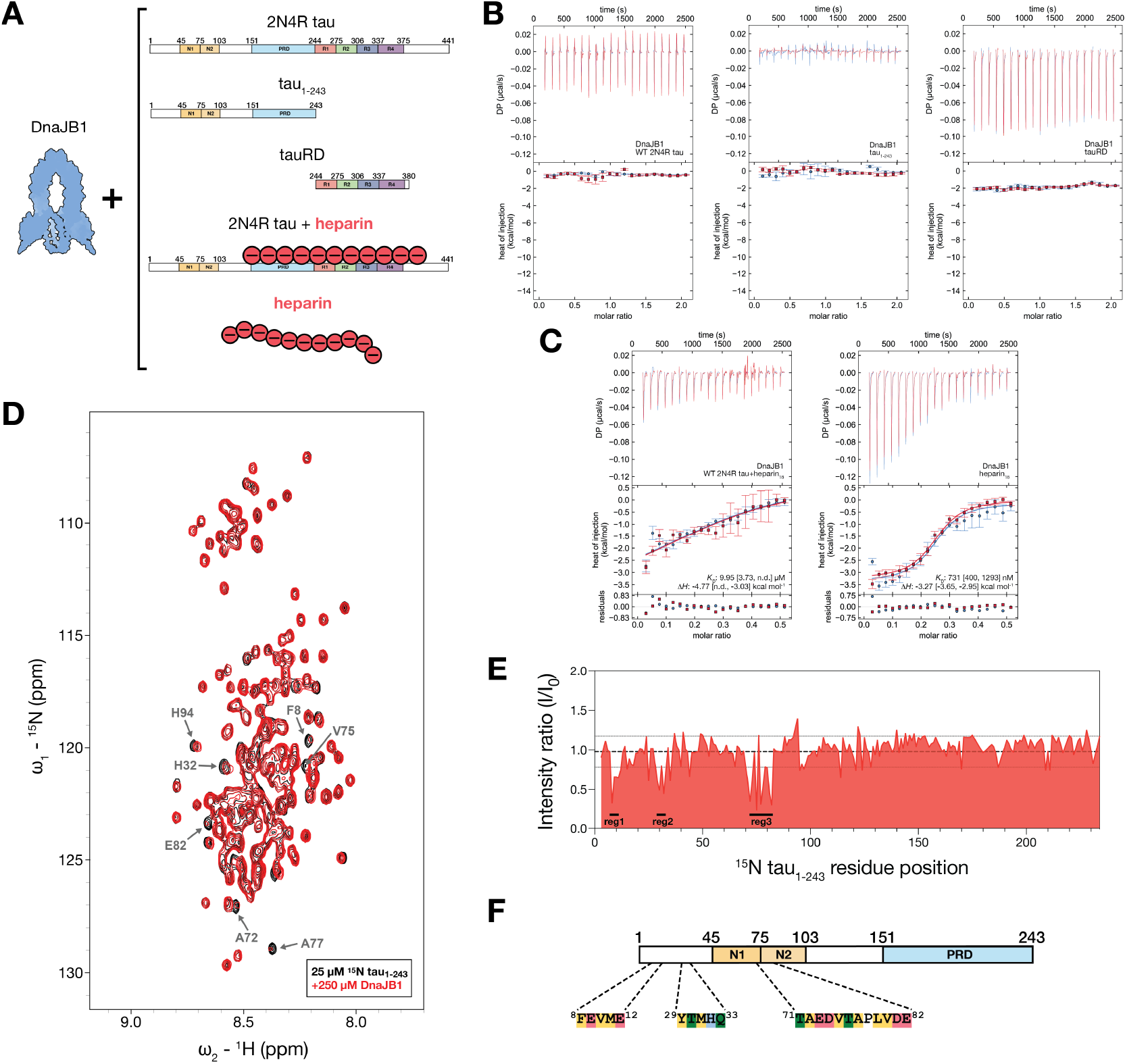
Weak DnaJB1:tau interactions are enhanced by the polyanionic factor heparin. **(A)** Schematic of protein constructs used in affinity measurement experiments. **(B**,**C)** ITC titrations of DnaJB1 binding to 2N4R WT tau, tau_1-243_, tauRD, 2N4R WT tau preincubated with equimolar concentration of heparin, and heparin alone. The top panels show reconstructed thermograms from NITPIC, the middle panels show binding isotherms and individual fits, and the bottom panels show the fitting residuals, when binding was detected. Error bars on reconstructed thermograms represent estimated peak integration errors from NITPIC. Square brackets represent a 68.3% confidence interval determined by error surface projection in global analysis of duplicate experiments in SEDPHAT. n.d. not determined. **(D)** _1_H–_15_N HSQC solution NMR spectra of _15_N-labeled tau_1-243_ alone (black) and in the presence of 10-fold molar excess of DnaJB1 (red) reveals changes of specific peaks (indicated by arrows). **(E)** Summary of the peak intensity ratio I/I_0_ of tau_1-243_ (I_0_) and tau_1-243_ with DnaJB1 (I) from panel (D). Thin dotted lines represent the range of single standard deviation from the average intensity ratio across all residues of tau_1-243_ (bold dashed line). **(F)** Amino acid sequences of three DnaJB1-binding sites on tau_1-243_ identified by HSQC NMR experiments from panels (D and E), shown in the context of the tau_1-243_ construct’s domain architecture.

Because of the exceptional stability of monomeric FL WT tau in solution, a large body of literature that describes molecular chaperones’ activity on tau aggregation utilized heparin to produce recombinant tau fibrils but did not investigate possible direct interactions of this polyanion with chaperones. We wondered whether presence of such highly negatively charged polymer in studies investigating these charge-dependent interactions might have impacted previous results. Therefore, we investigated the effect of heparin on DnaJB1:tau interactions and found that the chaperone can recognize heparin preincubated 2N4R tau monomer with micromolar affinity in contrast to no detected binding in the absence of heparin consistent with the heparin polymer bridging interactions between the chaperone and tau (Fig. 3BC) (Irwin et al., 2021). We additionally tested whether DnaJB1 alone binds to heparin. Titration of heparin with the chaperone revealed a high affinity interaction with a *K*_*D*_ of 731 nM (Fig. 3C). It has been previously reported that multivalency may be important for binding of this polyanion to tau (Fichou et al., 2019). When we tested heparin polymers of sizes canonically used to induce tau fibrilization, heparin_15/18_ (average molecular weights of 15 kDa and 18 kDa, respectively), we measured affinities in the nanomolar range, while low-molecular-weight heparin enoxaparin sodium showed much weaker binding to DnaJB1 (heparin_4.4_; Supplementary Fig. 4A). Presence of ThT-positive signal together with size-exclusion chromatography experiments of heparin-incubated DnaJB1 suggest formation of large complexes (Supplementary Fig. 2D and 4B). Intriguingly, we discovered that larger heparin species can bind not only the FL tau and tauRD as reported before, but also the N-terminus of tau alone, likely through binding to the partially basic proline-rich domain (Fig. 1A and Supplementary Fig. 4C) (Zhu et al., 2010; Fichou et al., 2019).

Similarly to our observations, previous quantification of affinities between monomeric aSyn and DnaJB1 using MST yielded no significant binding (Wentink et al., 2020). The authors turned to solution NMR to investigate the weak binding between DnaJB1 and aSyn, discovering a short acidic motif at the C-terminus of aSyn as the chaperone recognition site. Therefore, we decided to collect ^1^H–^15^N heteronuclear single quantum coherence spectroscopy (HSQC) spectra in an effort to detect the low affinity DnaJB1-binding sites on monomeric tau_1-243_. We produced uniformly ^15^N-labeled tau_1-243_ enabling complete assignment of the spectra to the sequence using previosuly published assignments (BMRB Entry 28065) (Ukmar-Godec et al., 2020). Titration of unlabeled DnaJB1 into ^15^N tau_1-243_ revealed broadening of a small subset of peaks (Fig. 3D). The peak broadening behavior across titration corresponds to the weak, transient binding, similarly as observed for aSyn (Supplementary Fig. 5AB) (Wentink et al., 2020). Residues of tau_1-243_ that indicate change to their local environment upon DnaJB1 addition group into three distinct regions: ^8^FEVME^12, 29^YTMHQ^33^, and ^71^TAEDVTAPLVDE^82^ (Fig. 3EF). These binding sites are primarily composed of negatively charged and hydrophobic residues and exhibit varied behavior during DnaJB1 titration with some obtained peaks within these sites shifting and broadening (F8, T30, A77) or predominantly broadening (A72, V75, E82) across the titration (Supplementary Fig. 5B). Therefore, we conclude that weak binding of DnaJB1 chaperone to the N-terminus of tau depends on short, predominantly acidic peptide sequences.

### DnaJB1 suppresses tau aggregation when seeded with heparin-induced FL tau fibrils but not with heparin-less tau fibrils

Next, we tested whether DnaJB1 acts differentially on distinct tau seed types in cell models of tau aggregation. We produced seeds FL tau seeds in the presence or absence of heparin, again leveraging the S320F mutation to enable heparin-free aggregation reactions (Fig. 4A). We found that the seed types have different efficiencies of seeding, with heparin-induced FL fibrils appearing more potent when compared to heparin-less seeds made of FL S320F tau or S320F tauRD (Supplementary Fig. 6A). Observed changes may be due to intrinsic differences in fibrilization capacity between the constructs. In DnaJB1 KD cells, we measured an increase in tau aggregation when cells were seeded with heparin-induced FL tau fibrils, but there was no significant change in aggregation when the biosensor cells were treated with heparin-less FL tau fibrils (Fig. 4B). This observation suggests that the heparin polyanion present on tau fibrils may play a role in recruiting cellular factors and is consistent with our previous results (Fig. 2EI). Importantly, none of the heparin controls used in this study induced spontaneous aggregation of tau in cells, therefore, all previously seen changes in tau inclusion formation likely arise from heparin-preincubated fibrils and not the heparin itself (Supplementary Fig. 6B).

**Figure 4.**
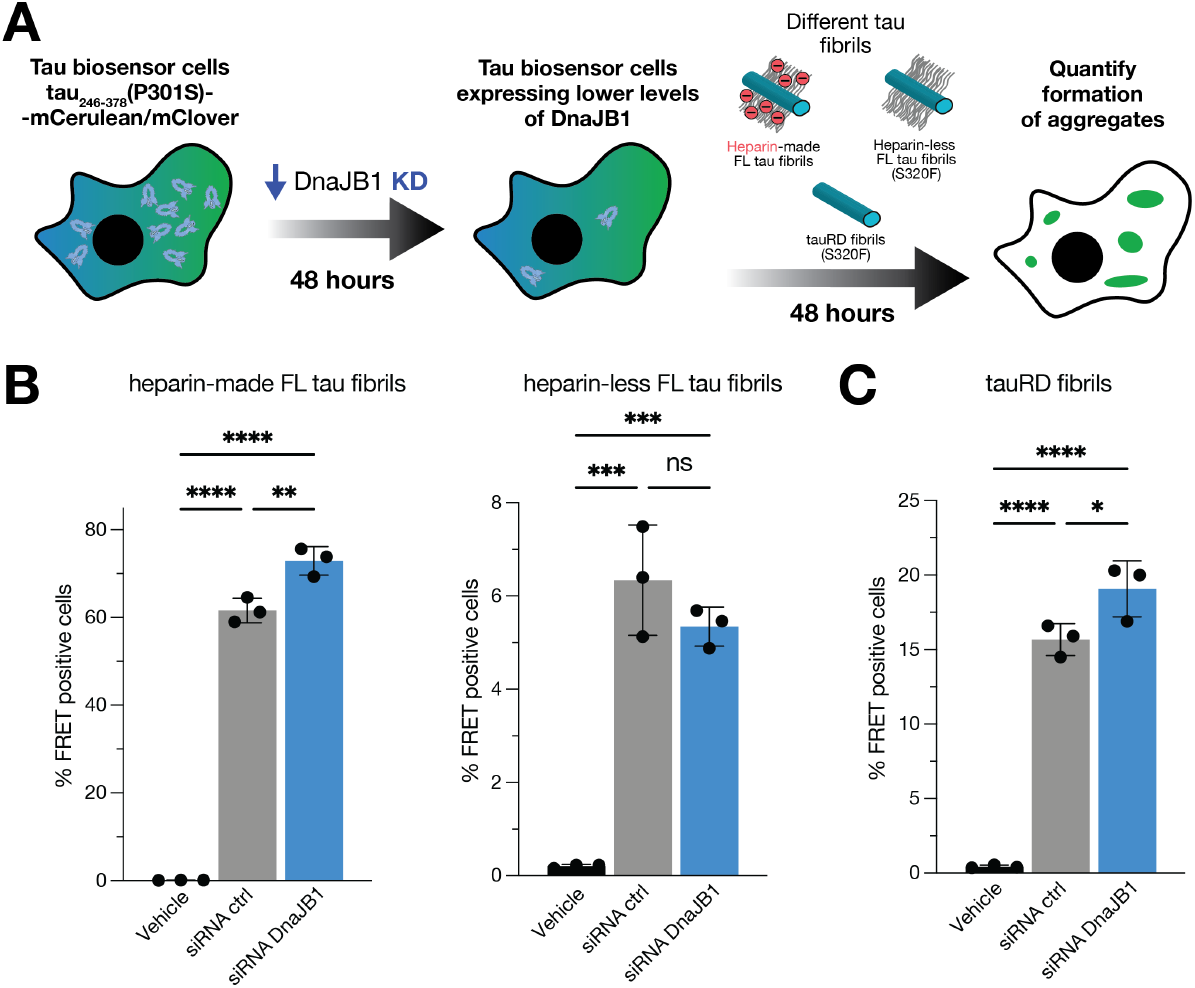
DnaJB1’s impact on tau aggregation depends on tau seed species. **(A)** Schematic of the experimental setup to test seeding of different tau constructs in biosensor cells stably expressing tauRD with either mCerulean3 or mClover3 fluorescent tag in the context of DnaJB1 knockdown. Tau aggregation was measured as a FRET signal via flow cytometry. **(B)** Lipofectamine-mediated tau seeding in tau biosensor cells using fibrils made with different tau constructs as seeds in the context of DnaJB1 knockdown. Cells were seeded with recombinant fibrils composed of heparin-induced FL 2N4R WT tau (left), heparin-free FL S320F tau (right). **(C)** Lipofectamine-mediated tau seeding in tau biosensor cells using fibrils made with heparin-free S320F tauRD lacking the N-terminal 243 residues, in the context of DnaJB1 knockdown. Data are shown as averages with error bars representing SD across 3 independent transfections. ns=not significant, (*) p < 0.05, (**) p < 0.01, (***) p < 0.001, (****) p < 0.0001

Subsequently, we investigated whether the presence of the N-terminal fuzzy coat in tau fibril seeds added to cell biosensors influences recognition by DnaJB1. We used two distinct heparin-less tau fibril types with the spontaneously aggregating S320F mutation: FL tau (residues 1-441) and tauRD (244-380), and found that, surprisingly, the removal of the N-terminal region of tau in seeds results in increased tau aggregation upon DnaJB1 knockdown (Fig. 4C). The observed role of DnaJB1 on seeding competency of the heparin-free tauRD fibrils is in agreement with the previous in vitro experiments that showed co-sedimentation of DnaJB1 chaperone with heparin-induced tauRD fibrils (Irwin et al. 2021).

We next sought to test whether DnaJB1 may influence aSyn seeded aggregation using a system analogous to the one used here for tau and DnaJB1 (Chlebowicz et al., 2023). Previously, in vitro studies showed the recognition and disassembly of aSyn fibrils by DnaJB1 as a part of Hsp70 machinery (Gao et al., 2015; Wentink et al., 2020). However, when DnaJB1 was knocked out using CRISPR in cellular models of aSyn aggregation, there was no effect on inclusion formation, in contrast to the strong suppression observed with DnaJB6 knockout (Deshayes et al., 2019). We hypothesized that the complete loss of DnaJB1 may have triggered compensatory activity from other JDPs, as functional redundancy between DnaJB1 and DnaJB4 has been reported (Nachman et al., 2020). To address this, we tested whether a partial knockdown of DnaJB1 alters aSyn seeding using the siRNA approach in previously developed A53T aSyn-CFP/YFP biosensor cells (Supplementary Fig. 6DF) (Chlebowicz et al., 2023). Upon lipofectamine delivery of recombinant FL WT aSyn fibrils into aSyn biosensors, we observed a 40% increase in aSyn aggregation in the DnaJB1 knockdown condition when compared to siRNA control (Supplementary Fig. 6E). Importantly, a recent study that utilized shRNA knockdown of DnaJB1 in SH-SY5Y cells reported a similar effect in inducing aSyn aggregation levels (Huang et al., 2025). Our findings suggest that reduction of DnaJB1 levels influences seeded aggregation in cellular models in a seed-dependent manner.

### DnaJB1’s JD and CTDI/CTDII are the tau-binding domains

To examine the surfaces of DnaJB1 that engage tau, we employed two different constructs: FL tau and the fuzzy coat-mimicking tau_1-243_. Interaction sites were identified by cross-linking mass spectrometry (XL-MS) with disuccinimidyl suberate (DSS) which modifies lysine residues to capture contact regions within the complex (Fig. 5A). This approach was successfully used to detect transient interactions between chaperones and their clients (Hou et al., 2021; Ayala Mariscal et al., 2022; Nitika et al., 2022). This cross-linker chemistry provides coverage of pertinent domains for both tau and DnaJB1, except for the G/F region of DnaJB1 which is depleted in lysines (Supplementary Fig. 7A) (Iglesias et al., 2009). First, we tested whether heparin can stabilize heterocomplex formation between DnaJB1 and tau as observed by ITC. Indeed, we found more prevalent DnaJB1:tau complex formation using SDS-PAGE after cross-linking (Supplementary Fig. 7B), as well as increased number of inter-protein cross-links relative to heparin-present conditions using the same statistical threshold (Fig. 5B). Analyzing the identified contacts, we found that the J-domain (JD) of DnaJB1 binds to the C-terminal end of the proline-rich domain (PRD) of FL tau in the absence of heparin with the results showing modified lysines K21 and K35 on and downstream of helix II of the JD (Fig. 5C and Supplementary Fig. 7C). In a recent cryo-EM study, thehe JD of DnaJB1 was found to participate in interactions with aSyn fibrils (Monistrol et al., 2025). In the presence of heparin, we found additional contacts between this region of DnaJB1 and the N-terminal boundary of PRD of tau (K148 and K150). Interestingly, we also detected contacts between K195 in the β-sandwich C-terminal domain I (CTDI) of DnaJB1 and PRD of tau when associated with heparin (Fig. 5C and Supplementary Fig. 7C). When we used the N-terminal tau_1-243_ as a client to mimic DnaJB1:fibril interactions, we observed an increase in detected DnaJB1:tau contacts (Fig. 5B and Supplementary Fig. 7C). In this case, both N- and C-terminal regions of tau’s PRD were bound by the chaperone independent of heparin. Previously described JD interaction sites with tau’s PRD were also enriched in cross-linked contacts. Newly identified contacts between CTDI of DnaJB1 and tau span domain of the chaperone, with additional interactions found at the ‘hinge’ between CTDI and CTDII (K242), as well as at K296 on CTDII of DnaJB1 in the presence of heparin (Fig. 5C and Supplementary Fig. 7C).

**Figure 5.**
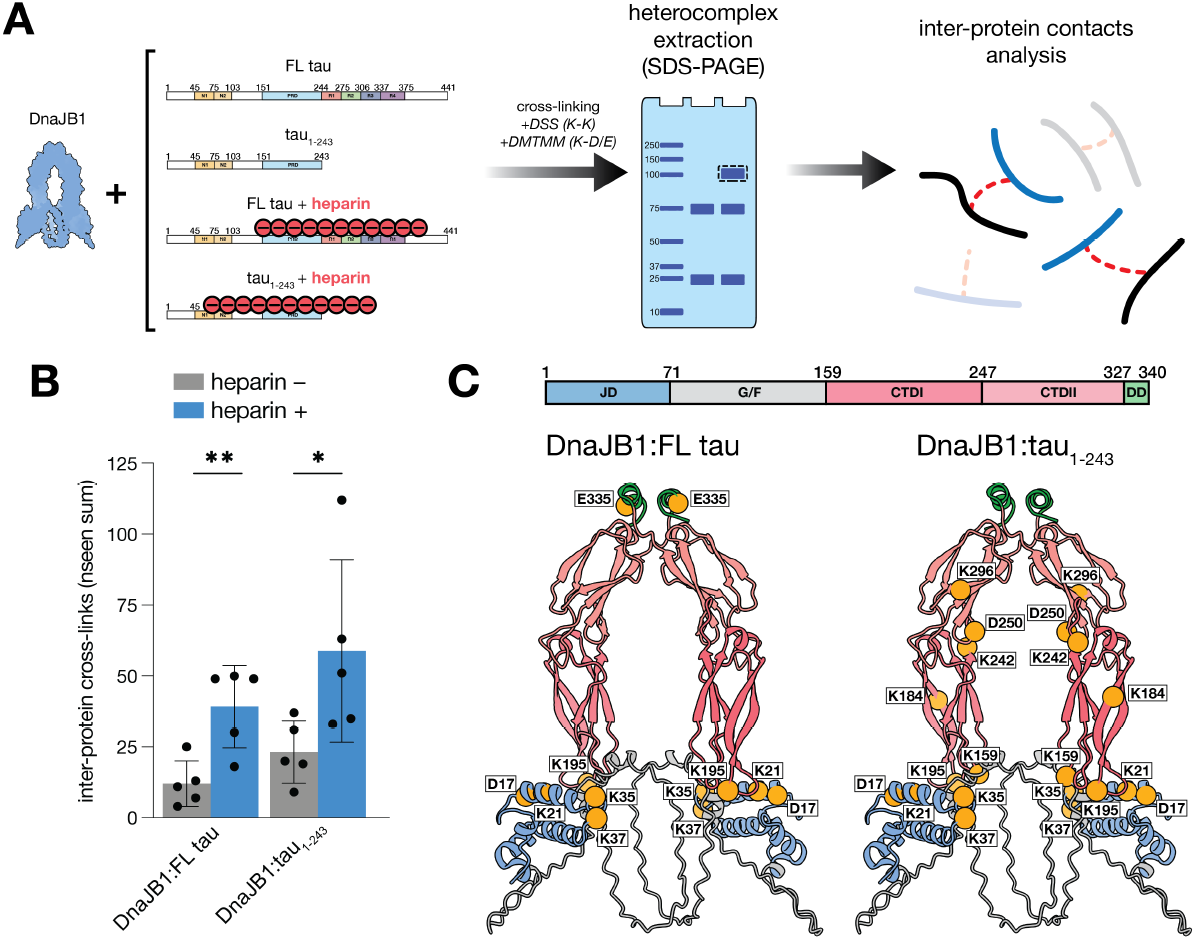
Client-recognition sites of DnaJB1 are clustered in its J-domain and the CTDI/CTDII interface. **(A)** Schematic of cross-linking mass spectrometry (XL-MS) experiments with isolation of the heterocomplex band from SDS-PAGE and subsequent analysis of the inter-protein contacts. **(B)** Quantification of the summarized inter-protein cross-link frequency in DSS datasets when the experiments were done in absence (gray) or presence (blue) of the polyanionic factor heparin. Data are shown as averages with error bars representing SD across 5 technical replicates. (*) p < 0.05, (**) p < 0.01. Symbols represent individual values across 5 technical replicates. **(C)** Identified tau-interacting sites mapped onto AlphaFold2 multimer model of the full-length human DnaJB1 chaperone in the form for a functional dimer. The coloring scheme of DnaJB1 domains from the N-to C-terminus: J-domain (JD, blue), glycine-phenylalanine rich linker (G/F, gray), C-terminal domains I (CTDI, dark red) and II (CTDII, pale red), dimerization domain (DD, green). Yellow spheres represent all tau-interaction sites identified using both tested cross-linker chemistries DSS (Lys-Lys) and DMTMM (Lys-Asp/Glu) in heterocomplexes of DnaJB1 with FL 2N4R tau (left) or tau_1-243_ (right).

The unstructured G/F region of DnaJB1 lacks lysine residues, which makes this region of the chaperone invisible to our DSS-based cross-linking studies. To complement the DSS experiments, we also used 4-(4,6-dimethoxy-1,3,5-triazin-2-yl)-4-methyl-morpholinium chloride (DMTMM) zero-length cross-linker that cross-links lysines with acidic residues (Leitner et al. 2014) (Supplementary Fig. 7A). We did not observe substantial contacts between tau and G/F region of DnaJB1, however, contacts to the JD as well as the boundary between CTDI/CTDII were confirmed with this alternative cross-linking chemistry (Fig. 5C and Supplementary Fig. 7D). We also identified contacts between the JD of DnaJB1 and the region upstream of PRD of tau (i.e. K132). Additionally, in the DnaJB1:FL tau dataset, we observed an inter-protein contact between one of tau’s amyloid motifs ^275^VQIINK^280^ and Dimerization Domain (DD) of DnaJB1. The DD together with CTDs were identified as client-interacting domains of DnaJB1 in previous NMR studies that used heparin-preincubated tau (Supplementary Fig. 7D) (Irwin et al., 2021).

To complement our ITC affinity measurements between DnaJB1 and heparin, we also leveraged information from our XL-MS experiments to deduce potential heparin-interacting regions on this chaperone by quantifying differences in the frequency of DSS chemical modification at lysine residues as a proxy for solvent accessibility (Supplementary Fig. 7E). We observed a heparin-dependent decrease in mono-link frequency at K286 when FL tau and tau_1-243_ were used as clients, suggesting that this site near CTDII may be involved in binding to heparin. Importantly, when we compared the charge distribution of DnaJB1 in this region, we found a cluster of basic residues that could be compatible with heparin binding (Supplementary Fig. 7F).

## Discussion

Aggregation of misfolded proteins into pathological amyloid fibrils causes significant disruption in the protein homeostasis network (Hipp et al., 2014). JDPs are critical members of this network and directly bind misfolded clients recruiting them to the Hsp70 chaperone machinery for subsequent ATP-dependent processing. Unraveling the biochemical and structural details of how chaperones recognize pathology-associated misfolded proteins remains a key goal with a great therapeutic potential. In this study, we describe how a member of JDP family, molecular chaperone DnaJB1, binds to tau protein whose misfolding is linked to numerous neurodegenerative diseases. We found that in the cellular model of tau aggregation the decreased expression of this chaperone leads to enhanced tau aggregation efficiency when the N-terminal acidic region of this protein remains accessible. These findings shed light on the prospective chaperone-recruiting role of the previously understudied N-terminal region of tau which remains disordered in tau fibrils and forms a characteristic fuzzy coat along the fibril. In this fuzzy coat, distinct acidic residue-containing motifs are recognized by DnaJB1, similarly to what has been postulated for aSyn fibrils where DnaJB1 recognizes a short peptide sequence mainly composed of negatively charged and hydrophobic residues located on the disordered C-terminal tail (Wentink et al., 2020; Jäger et al., 2024). Despite the fact that tau is an intrinsically disordered protein, it is thought that FL tau monomers are engaged in long-range intra-molecular interactions between the acidic N-terminus and basic central region of the protein that includes its repeat domain (Jeganathan et al., 2006, 2008; Hou et al., 2021; Marien et al., 2025). This conformation may play a protective role while also prohibiting DnaJB1 from binding to the acidic part of tau (Fig. 6). Heparin, a polyanionic factor previously linked to tau pathology and used to induce tau fibrilization in vitro, binds with high affinity to tau’s basic region. This disrupts intra-molecular electrostatic interactions, exposing the N-terminal region for DnaJB1 binding, which may be facilitated by heparin through interactions between basic residues of both proteins (Sibille et al., 2006; Zhu et al., 2010; Zhao et al., 2017). The fuzzy coat of tau fibrils has also been proposed to change fibril’s mechanical properties through the interactions of the N-terminal acidic region with basic surfaces of the fibril core and thus also altering its accessibility to cellular factors (Fig. 1A) (Wegmann et al., 2013). Co-localization of heparan sulfate and neurofibrillary tangles in patient tissues points at its potential importance in pathological amyloid fibrils (Goedert et al., 1996). Similarly to tau monomers, presence of this polyanionic factor may change the conformation of the N-terminus of tau now enabling DnaJB1 to bind to the negatively charged residues of the fuzzy coat, which may be also applicable to other fibril-interacting proteins in the cellular context.

**Figure 6.**
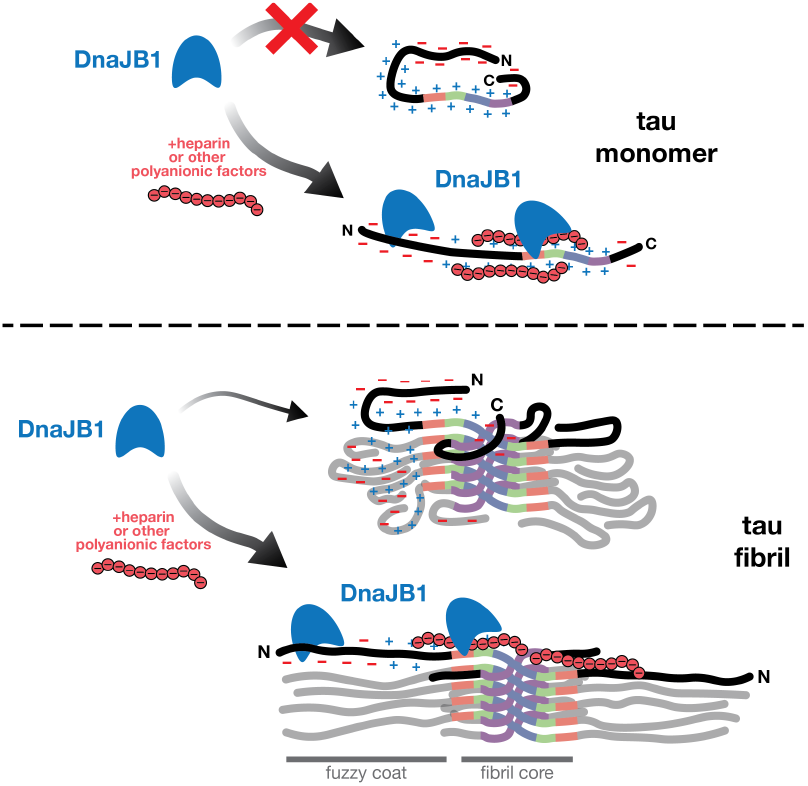
Speculative model of interactions between DnaJB1 chaperone and different tau species. **(Top)** In monomeric form, 2N4R tau protein adopts conformations resembling the ‘paperclip’ model with its acidic N- and C-termini engaged in transient contacts with the basic core of the protein. Upon binding of the polyanionic factor heparin, the tau monomer now adopts more open conformation with the acidic N-terminus now accessible to be recognized and bound by DnaJB1 chaperone, which can also recognize the anionic charge of heparin that coats the basic proline-rich and repeat domains of tau. **(Bottom)** Similarly to the monomeric form, in the absence of heparin the fuzzy coat of 2N4R tau fibrils is more compacted with acidic and basic regions of the N-terminus of tau transiently interacting with each other. However, the presence of heparin disengages these contacts and now accessible acidic regions of the fuzzy coat may be bound by DnaJB1.

It has been reported that DnaJB1 uses its C-terminal domains to bind clients, and in this study, we report similar domain utilization of this chaperone (Suzuki et al., 2010; Irwin et al., 2021; Ayala Mariscal et al., 2022). We also identified the conserved JD to play a role in tau recruitment. The canonical role of the JD is to mediate binding to Hsp70 and its ATPase activity, however, recent data provide a new perspective of this region as a potential client-binding site, as observed recently for aSyn fibrils (Kampinga & Craig, 2010; Rosenzweig et al., 2019; Monistrol et al., 2025). Client-interaction sites located at helix II of JD suggested by our results present a novel interpretation as this domain encodes capacity to recruit Hsp70 to clients for downstream processing. However, Hsp70-JDP interaction may be inhibited by helix V on the G/F region of DnaJB1 (Faust et al., 2020). Our findings suggest additional, client-binding role of this region of DnaJB1’s JD and further studies should explore the role of substrate-binding events at this site in the context of autoinhibition and Hsp70-binding.

Our findings suggest that the weak, transient binding of DnaJB1 to tau becomes enhanced in the presence of heparin. The interaction of heparin-related sulfated polysaccharides with tau has been extensively studied, however, their potential influence on the protein homeostasis network, in particular chaperones, has not been yet investigated. Moreover, the relation of chaperones to the numerous other polyanionic factors with important cell functions remains to be reported. For example, polyphosphates found to be at micromolar concentrations in the cells, modulate aggregation of several proteins into amyloid fibrils, including tau and α-synuclein (Cremers et al., 2016; Barolo et al., 2025). Therefore, we underscore the possibility of a potential bias of previous in-cell proteomics-based studies of tau aggregation that may have been caused by presence of heparin. Our data suggest that this polyanionic factor may alter the interactome of proteins binding to aggregates.

Studies investigating chaperone binding to tau protein were often limited by utilization of heparin to induce tau fibrilization, and/or using tau constructs that contain only the basic repeat domain. It has been established that heparin-induced tau fibrils are structurally different from any fibrils isolated from patient samples questioning the physiological relevance of this factor in studies of tau aggregation (Zhang et al., 2019). In studies of aggregation prevention by cellular factors, the observed effect may not reflect direct chaperone protection of soluble tau from adopting β-sheet conformations, but rather the depletion of heparin, as it binds to charged patches on chaperones and thereby reduces its fibril-inducing activity. Our results show that DnaJB1 prevents formation of tau seeds only in the absence of heparin, suggesting that previous in vitro studies of DnaJB1:tau interactions may have relied on residual heparin bridging chaperone–fibril interactions. By including the N-terminal region of tau, our experiments expand the potential chaperone-binding sites on this aggregation-prone protein. Importantly, the solvent-exposed N-terminal fuzzy coat of tau fibrils can also be recognized by other cellular components, making it an appealing target for therapeutic strategies such as fibril–specific antibody design (Wentink et al., 2020; Jäger et al., 2024; Parlato et al., 2025). DnaJB1 binding to the fibril’s disordered region of aSyn recruits Hsp70 and is an important pre-requisite of the entropic-pulling model of amyloid disaggregation (Wentink et al., 2020; Monistrol et al., 2025).

Given the complexity of the Hsp70 chaperone machinery, with Hsp70 playing the central role with the help of co-chaperones, JDPs and nucleotide-exchange factors, future studies need to address how addition of these components influences findings presented in this study. There are more than 10 members of Hsp70s family and more than 40 other JDPs expressed in human cells that may possibly interact differently with FL tau fibrils with and without the presence of heparin. Our results show very limited effect on tau aggregation when DnaJB1’s expression levels were manipulated in cell models. Experiments that change expression levels of DnaJB1 chaperone with parallel investigation of the role of other B class members, such as DnaJB2 or DnaJB4 that has been shown to bind a similar client pool, are necessary to study the interplay and potential degeneracy in the action of this co-chaperone family (H.-J. Chen et al., 2016; Nachman et al., 2020; Inoue et al., 2024). Moreover, it will be important to test whether the composition and action of the protein quality control systems and tau’s interactome significantly change when more pathology-relevant models of tau aggregation are utilized, such as iPSC-derived neuronal FL tau biosensors. The potential role of heparin-related factors has been postulated in the context of Alzheimer’s disease, but impact, if any, of such large and highly charged molecules on the intrinsic protein homeostasis network and their clients has not yet been deconvoluted (Goedert et al., 1996). Future studies should also investigate how charge-changing post-translational modifications of tau influence recognition by chaperones. The charge distribution change of tau monomer that is caused by the pathology-related phosphorylation may potentially lead to: (1) change of conformation into more templating-competent that directs tau towards aggregation, as well as (2) differential recruitment of binding partners whose interactions depend on salt bridges, as has been observed for chaperones. Importantly, the preference of DnaJB1 to bind phosphorylated protein has been recently reported for α-synuclein fibrils (Pacheco et al., 2024).

There is still no consensus on how chaperones recognize amyloid fibrils, with two non-mutually exclusive strategies. One view suggests interaction at the open ends of the long fibrils where it may be less energetically demanding to break off individual monomers from the tip of the fibrils by depolymerization (Franco et al., 2021; Schneider et al., 2021; Beton et al., 2022). The alternative view is that these interactions may occur along the fibril axis with the present fuzzy coat creating abundant binding sites, which increases the probability of clustering chaperones to deform fibrils thus leading to their fragmentation (Gao et al., 2015; Wentink et al., 2020; Jäger et al., 2024). Consistent with this idea, DnaJB1’s low affinity for the N-terminus of tau in tau monomers would be enhanced through avidity to this element in fibrils. Obtaining a high-resolution structure of DnaJB1 (as well as the whole Hsp70 machinery) in complex with tau in monomeric and/or fibril form will be particularly challenging due to the transient nature of the chaperone– client interactions and heterogeneity on fibril surfaces. Nevertheless, new studies have taken advantage of the latest developments in atomic force microscopy, cryo-EM, and cryo-ET methods to fill this gap, in particular in the context of aSyn fibrils (Franco et al., 2021; Beton et al., 2022; Monistrol et al., 2025). These exciting next studies will provide the invaluable biochemical and structural insight into the interactions between the cellular machinery and the neurodegeneration-associated clients to inform development of new therapeutic interventions.

## Materials and methods

### Cell biosensors

#### DnaJB1 overexpression/knockdown and induction of tau aggregation in biosensor cells using N-terminal tau truncations with mEOS3.2

The constructs containing N-terminal truncations of tau (longest construct: residues 1-408) with C-terminal mEOS3.2 fluorescent tag were made using previously obtained YFP tagged versions of the same truncations in FM5 backbone with digestion with BamHI and AscI restriction enzymes. Twelve total constructs were obtained with ∼20 residue step truncations starting from the N-terminus of tau protein (therefore the list of truncations tested with residue numbers referring to the human 2N4R isoform of tau: 1-408, 22-408, 42-408, 62-408, 82-408, 103-408, 122-408, 142-408, 162-408, 182-408, 222-408, 242-408 with omission of 202-408 truncation due to difficulties in obtaining this construct through cloning. Obtained constructs were confirmed with Sanger sequencing. mEOS3.2 fluorescent tag has been successfully used for studying aggregation-prone proteins and protein interactions, however future studies should consider alternative tagging or tag-free approaches to further validate findings in this manuscript.

HEK293T cells were plated in 96-well plates at 2,000 cells/well in 130 μL of high glucose DMEM (supplemented with 10% (v/v) FBS, 1X GlutaMAX, and penicillin-streptomycin) in three technical replicates. 24 hours later, the cells were treated with 10 μL total volume of either FM5 plasmid with DnaJB1-mCerulean3 under UbC promoter (or only mCerulean3 as a control), or siRNA oligo duplex targeting DnaJB1 (or scrambled control) with sequences as indicated below:

siDnaJB1 (Origene #SR320570C): GCAUGGACAUUGAUGACCCAUUCTC siControl (Origene #SR30004): CGUUAAUCGCGUAUAAUACGCGUAT Scrambled siRNA was used as a control to account for potential off-target effects of the siRNA approach. For overexpression conditions 49.5 ng of DNA/well was used together with 0.25 μL/well of Lipofectamine 2000 (Invitrogen #11668500). For knockdown conditions 50 nM final concentration of siRNA duplex was used together with 0.421 μL/well of Lipofectamine RNAiMAX reagent (Invitrogen #13778075). In OE as well as KD conditions after 48 hours, DNA encoding N-terminal tau truncations with mEOS3.2 fluorescent tag were added at 49.5 ng of DNA/well together with 0.25 μL/well of Lipofectamine 2000 in 10 μL total volume. OE and KD were confirmed to be effective after 48 hours, and this timepoint has been selected allowing for maintenance of the changed chaperone expression level for the duration of the experiment. After 48 hours, media were replaced, and 100 nM final concentration (monomer concentration) of heparin-induced recombinant 2N4R tau fibrils was added together with 0.75 μL/well of Lipofectamine 2000 in 20 μL total volume. Prior to cell treatment, the tau fibrils were sonicated for 30 s at an amplitude of 65 on a Q700 Sonicator (QSonica) that has a maximum power output of 700 watts. 48 hours after treatment with tau fibrils, the cells were harvested by 0.05% trypsin digestion and then fixed in 1X Dulbecco’s PBS (DPBS) with 2% paraformaldehyde (PFA) for 10 minutes at room temperature. The cells were then washed and resuspended in 150 μL of 1X DPBS. This set of experiments was done in 3 biological replicates (3 technical replicates each) defined as each replicate separated by at least 2 cell passages.

#### DnaJB1 overexpression/knockdown and induction of tau aggregation in cells using tauRD biosensor cell lines and seeding with different tau fibrils

HEK293T tauRD P301S v2L FRET biosensor cell line that stably expresses mClover3 or mCerulean3 tagged tauRD (residues 246-378) monomer was obtained from the Diamond lab (Hitt et al., 2021). Cells were plated in 96-well plates at 17,000 cells/well in 140 μL of media. 24 hours later, the cells were treated with DNA (DnaJB1 overexpression) or siRNA duplexes (DnaJB1 knockdown) as described above. After 48 hours, media were replaced with a fresh 130 μL, and fibrils made of three different tau constructs were added at 100 nM final concentration (monomer concentration) together with 0.75 μL/well of Lipofectamine 2000 in 20 μL total volume, with prior sonication as described above. 48 hours after treatment with tau fibrils, cells were harvested and fixed for flow cytometry, as described above. These experiments were done as three independent transfections.

#### DnaJB1 overexpression/knockdown and induction of α-synuclein aggregation using α-synuclein biosensor cell lines

HEK293T biosensor cells stably expressing monomeric α-synuclein (A53T) with either CFP or YFP fluorescent tag were obtained from the Diamond lab (Chlebowicz et al., 2023). Cells were plated in 96-well plates at 17,000 cells/well in 140 μL of complete DMEM. 24 hours later, the cells were treated with DNA (DnaJB1 OE) or siRNA duplexes (DnaJB1 KD) as described above. After 48 hours, media was replaced with a fresh 130 μL, and recombinant fibrils made of WT α-synuclein at 100 nM final concentration (monomer concentration) together with 0.75 μL/well of Lipofectamine 2000 in 20 μL total volume. Prior to cell treatment, α-synuclein fibrils were sonicated at an amplitude of 65 for 5 min (1 min ON and 1 min OFF) using a Q700 Sonicator (QSonica). 48 hours after treatment with α-synuclein fibrils, cells were harvested and fixed for flow cytometry, as described above. These experiments were done as three independent transfections.

### Flow cytometry

#### Flow cytometry of cells expressing mEOS3.2-tagged N-terminal tau truncations

Invitrogen Attune CytPix flow cytometer was used to perform FRET signal measurements. Prior to data collection, mEOS3.2 in cells was photoconverted with LED UV lamp (Chauvet) for 20 minutes at 12W power at a distance of approximately 5 cm from the top of the plate. To measure mCerulean3 signal, cells were excited with 405 nm laser, and 440/50 nm filter was used. To measure non-photoconverted (green) mEOS3.2 and FRET signal (between non-photoconverted (green) mEOS3.2 and photoconverted (red) mEOS3.2), cells were excited with 488 nm laser and filters 530/30 and 610/20 were used, respectively. To measure photoconverted (red) mEOS3.2 signal, 561 nm laser was used, with 620/15 nm filter. To quantify FRET, gating was conducted as established previously (Holmes et al., 2014; Furman et al., 2015) and shown at Supplementary Fig. 1C, where additional gating was done for overexpression conditions with mCerulean3 as the reporter of DnaJB1. Data analysis was done using FlowJo 10.10.0 (BD) and the results are reported as percentage changes in FRET-positive cells percentage (of parent population) of DnaJB1-mCerulean3 vs. mCerulean3 (for DnaJB1 overexpression) or siDnaJB1 vs. siControl (for DnaJB1 knockdown).

#### Flow cytometry of tauRD biosensor cells

Invitrogen Attune NxT CytKick flow cytometer was used to perform FRET signal measurements. To measure mCerulean3 and FRET signal cells were excited with 405 nm laser and filters 440/50 nm and 512/25 nm were used, respectively. To measure mClover3 signal, 488 nm laser was used for excitation, and filter 530/30 nm was applied. To quantify FRET, gating was conducted as established previously (Holmes et al., 2014; Furman et al., 2015) and shown at Supplementary Fig. 6C. Data analysis was done using FlowJo 10.10.0 (BD). For testing whether DnaJB1 knockdown changes tau assembly efficiency after treatment with different tau fibrils (Fig. 4BC) a one-way analysis of variance with Tukey’s correction for multiple comparisons was used to assess statistical significances relative to Lipofectamine vehicle and siRNA controls in GraphPad Prism 10.2.3. For testing efficiencies of different tau fibrils in inducing tau assembly (Supplementary Fig. 6A) a one-way analysis of variance with Dunnett’s correction for multiple comparisons was used to assess statistical significance relative to Lipofectamine vehicle control. For testing whether heparin alone can induce tau assembly (Supplementary Fig. 6B) a one-way analysis of variance with Tukey’s correction for multiple comparisons was used to assess statistical significances relative to no treatment and 1X DPBS controls.

#### Flow cytometry of α-synuclein biosensor cells

Invitrogen Attune NxT CytKick flow cytometer was used to perform FRET signal measurements. To measure CFP and FRET signal cells were excited with 405 nm laser and filters 440/50 nm and 512/25 nm were used, respectively. To measure YFP signal, 488 nm laser was used for excitation, and filter 530/30 nm was used. To quantify FRET, gating was conducted as established previously (Holmes et al., 2014; Furman et al., 2015) and shown at Supplementary Fig. 6F. Data analysis was done using FlowJo 10.10.0 (BD). For testing whether DnaJB1 knockdown changes α-synuclein assembly (Supplementary Fig. 6E) a one-way analysis of variance with Tukey’s correction for multiple comparisons was used to assess statistical significances relative to Lipofectamine vehicle and siRNA controls in GraphPad Prism 10.2.3.

### Immunoblotting

Immunoblotting of the chaperone and tau was conducted as established before (Perez et al., 2023). HEK293T cell pellets were lysed using RIPA buffer (Thermo Scientific) containing one tablet of cOmplete mini EDTA-free protease inhibitor (Roche) per 10 mL buffer by trituration with insulin syringes and subsequent 15 minutes incubation on ice. Cell debris was removed by centrifugation at a speed of 17,200 x g at 4 °C for 15 minutes, and supernatants were transferred to fresh tubes and protein concentration was measured using Pierce BCA Protein Assay Kit (Thermo Fisher).

5 μg of total protein samples were prepared using 4X Bolt LDS Sample Buffer (Invitrogen) supplemented with freshly added 10% (v/v) 2-mercaptoethanol (BME) and then heated at 98°C for 5 minutes. SDS-PAGE using NuPAGE Bis-Tris Midi Protein Gels 4-12% (Invitrogen) was conducted to resolve proteins with subsequent transfer onto Immobilon-P PVDF membranes (Millipore Sigma) using a Trans-blot semi-dry transfer cell (Bio-Rad). The membranes were blocked in 1X TBST buffer (25 mM Tris, 137 mM NaCl, 2.7 mM KCl, 0.1% (v/v) Tween-20, pH 7.4) containing 5% (w/v) non-fat milk powder (Bio-Rad). All incubations of the membranes with antibodies were done using 1X TBST buffer with 5% (w/v) milk powder.

Antibodies used for immunoblotting in this study are as follows, with primary: rabbit polyclonal anti-DnaJB1 (Cell Signaling #4868) at 1:1,000 dilution; mouse monoclonal anti-tau MD2.1, developed as a part of previous study (Hitt et al., 2023), at 1:10,000 dilution; mouse monoclonal anti-β-actin (Proteintech #66009-1) at 1:5,000 dilution; and secondary: donkey anti-rabbit Amersham ECL HRP-linked F(ab’)_2_ fragment (Cytiva #NA9340) at 1:10,000 dilution; goat anti-mouse H&L (HRP) (Abcam #ab6789) at 1:10,000 dilution.

Quantification of band intensities was done using Fiji distribution of ImageJ 2.14.0 (Schindelin et al., 2012) with background subtraction from each DnaJB1, tau, and β-actin band. The signals for all conditions were normalized to the signal of the Lipofectamine vehicle control. Replicate 2 from mCerulean3 condition has been omitted due to sample loss during lysate preparation. Final annotated figures were composed using LabFigures tool available at sciugo.com. Raw uncropped pictures of membranes and intensity measurements are available in the Source Data file.

### Protein expression and purification

Constructs with wild-type and point mutants (A152T, R406W) of the full-length 2N4R tau with C-terminal 6xHis tag in pET28b plasmid were used as published previously (D. Chen et al., 2019). 2N4R tau with S320F point mutation was used without 6xHis tag. All FL tau constructs were grown using E. coli BL21(DE3) cells in 2X YT medium at 37 °C with shaking 220 RPM until OD600 reached 0.8 and then induced with 1 mM final concentration of isopropyl β-d-1-thiogalactopyranoside (IPTG) for 4 hours at the same temperature and shaking speed. After spinning, cell pellet was resuspended in the lysis buffer (50 mM MES, 10 mM EDTA, 0.1 mM PMSF, 10 mM DTT, pH 6.0) supplemented with cOmplete mini EDTA-free protease inhibitor (Roche) and stored at -80 °C. Cells were lysed by addition of lysozyme at final concentration of 100 mg/mL of cell lysate, and subsequent sonication on ice for two rounds of 10 minutes each at 50% power and 50 pulse using Omni Sonic Ruptor 400 (Omni International). After clarification of the cell lysate, supernatant was filtered with 0.22 μm membrane filter in a vacuum system and loaded onto pre-equilibrated HiTrap SP HP 5 mL ion exchange chromatography column (Cytiva). For this run, a gradient of low salt (50 mM MES, 10 mM DTT, pH 6.0) and high salt (50 mM MES, 1 M NaCl, 10 mM DTT, pH 6.0) buffers was used, fractions containing construct of interest were identified by SDS-PAGE, pooled together and dialyzed overnight into 1X PBS, 20 mM imidazole, 1 mM BME at 4 °C. Dialyzed protein was loaded onto Ni-NTA agarose beads and eluted with 1X PBS, 500 mM imidazole, 1 mM BME. Eluted sample was concentrated using 10 kDa MWCO Amicon Ultra centrifugal filters (Millipore Sigma) and injected onto pre-equilibrated HiLoad 16/600 Superdex 200 size exclusion chromatography column (GE) and subsequently resolved in 1X PBS, 10 mM DTT, pH 7.4. SDS-PAGE was used to identify fractions containing purified protein, then protein was aliquoted, flash-frozen in liquid nitrogen, and stored at -80 °C until further use. For construct containing S320F point mutation Ni-NTA purification step was omitted.

Construct containing tau_1-243_ was used with C-terminal 6xHis tag in pET28b plasmid and was a kind gift from Zhiqiang Hou and Ayde Mendoza-Oliva. This construct was grown similarly as the FL tau construct, however TB medium was used instead of 2X YT and harvested cells were resuspended in the lysis buffer of 50 mM Tris, 500 mM NaCl, 1 mM BME, 20 mM imidazole, 1 mM PMSF, pH 7.0 supplemented with cOmplete mini EDTA-free protease inhibitor (Roche). After the cell lysis as described above, the clarified supernatant was loaded onto Ni-NTA agarose beads and eluted with 50 mM Tris, 250 mM NaCl, 1 mM BME, 500 mM imidazole, pH 7.0. The eluted sample was dialyzed overnight into 50 mM Tris, 50 mM NaCl, 1 mM BME, 10 mM DTT, pH 7.0 that was also the low-salt buffer during the subsequent ion exchange chromatography with HiTrap Q HP 5 mL column (Cytiva). Protein was eluted with gradient of low salt and high salt (50 mM Tris, 1 M NaCl, 1 mM BME, 10 mM DTT, pH 7.0) buffers, and SDS-PAGE was used to identify fractions containing purified protein, then protein was aliquoted, flash-frozen in liquid nitrogen, and stored at -80 °C until further use.

To produce ^15^N-labeled tau_1-243_ a single colony from a freshly transformed plasmid into BL21(DE3) cells was inoculated into 10 mL 1X LB medium and grown for 7-8 hours at 37 °C while shaking at 220 RPM. This 10 mL culture was then mixed into 100 mL of M9 medium (42.2 mM Na_2_HPO_4_, 22 mM KH_2_PO_4_, 8.5 mM NaCl, 0.1 mM CaCl_2_, 2 mM MgSO_4_, 0.05% (w/v) thiamine, 1X trace metals, 0.4% (w/v) glucose, and 0.1% (w/v) ^15^N-labeled NH_4_Cl) and grown overnight at 37 °C while shaking at 220 RPM. Then, cells were spun at 2,000 x g for 10 minutes, and the cell pellet was resuspended in 1 liter of M9 medium and grown until OD600 reached 0.6-0.8, and protein expression was induced by adding 1 mM IPTG for 4 hours at the same temperature and shaking speed. Cell lysis and further purification was conducted in the same manner as for the non-labeled protein described above.

The construct containing WT or S320F tauRD (residues 244-380) in pET29b with no tags was expressed in BL21(DE3) cells as described for FL constructs, however TB medium was used instead of 2X YT and harvested cells were resuspended in the lysis buffer of 20 mM MES, 1 mM EDTA, 1 mM MgCl_2_, 1 mM PMSF, 5 mM BME, pH 6.8 supplemented with cOmplete mini EDTA-free protease inhibitor (Roche). After the cell lysis with lysozyme and sonication as described above, the NaCl concentration was increased to 500 mM final concentration, the lysate was boiled for 20 minutes, and after spinning at 15,000 x g for 20 minutes, the supernatant was dialyzed overnight into 20 mM MES, 50 mM NaCl, 5 mM BME, pH 6.8 which was used as the low salt buffer for subsequent ion exchange chromatography with HiTrap SP HP 5 mL column. Protein was eluted with gradient of low salt and high salt (20 mM MES, 600 mM NaCl, 5 mM BME, pH 6.8) buffers, and SDS-PAGE was used to identify tauRD-containing fractions, that were subsequently pooled together, concentrated using 3 kDa MWCO Amicon Ultra centrifugal filters (Millipore Sigma) and injected onto pre-equilibrated HiLoad 16/600 Superdex 75 size exclusion chromatography column (GE) and eluted into 1X PBS, 10 mM DTT, pH 7.4. SDS-PAGE was used to identify fractions containing purified protein, then protein was aliquoted, flash-frozen in liquid nitrogen, and stored at -80 °C until further use.

The construct containing FL WT α-synuclein in pET28b with no tags was a kind gift from the Diamond lab and was purified as established previously (Beton et al., 2022). Protein was expressed in BL21(DE3) cells grown in LB Miller’s modified medium at 37 °C with shaking 220 RPM until OD600 reached 0.8 and then induced with 0.5 mM IPTG for 3 hours at the same temperature and shaking speed. After harvest, cell pellet was resuspended in the lysis buffer (100 mM Tris, 10 mM EDTA, 2 mM DTT, pH 8.0) supplemented with cOmplete mini EDTA-free protease inhibitor (Roche) and stored at -80 °C. Cells were lysed using probe sonication on ice for three rounds of 10 minutes each at 50% power and 50 pulse using Omni Sonic Ruptor 400 (Omni International) and clarified by spinning at 15,000 x g for 1 hour. The obtained supernatant was boiled for 30 minutes and spun down at 15,000 x g for 40 minutes. The resulting supernatant was supplemented with streptomycin sulfate in concentration of 30 mg per 1 mL of supernatant, and nutated for 1 hour at 4 °C. After centrifugation at 15,000 x g for 30 minutes, the supernatant was supplemented with ammonium sulfate at a concentration of 400 mg per 1 mL of supernatant, and nutated for 1 hour at 4 °C. After centrifugation, pellet containing α-synuclein protein was resuspended in 10 mL of 1X TBS (25 mM Tris, 137 mM NaCl, 2.7 mM KCl, pH 7.4) and dialyzed overnight at room temperature into 25 mM Tris pH 7.5. Obtained solution was loaded onto pre-equilibrated HiTrap Q HP 5 mL column (Cytiva), and a gradient of low salt (25 mM Tris, pH 7.5) and high salt buffers (25 mM Tris, 1 M NaCl, pH 7.5) was used for elution. Fractions containing purified protein were identified with SDS-PAGE, then protein was aliquoted, flash-frozen in liquid nitrogen, and stored at -80 °C until further use.

Plasmid containing full-length human DnaJB1 with N-terminal SUMO protein and 6xHis tag in pET29b was obtained from GenScript. Construct was expressed in BL21(DE3) cells grown in LB Miller’s modified medium at 37 °C with shaking 220 RPM until OD600 reached 0.8 and then induced with 1 mM IPTG overnight at 25 °C with 150 RPM shaking. After harvest, cell pellet was resuspended in the lysis buffer (50 mM HEPES, 500 mM KCl, 20 mM imidazole, 1 mM BME, 1 mM PMSF, pH 7.5) supplemented with cOmplete mini EDTA-free protease inhibitor (Roche) and stored at -80 °C. Cells were lysed with lysozyme and probe sonication on ice as described above, and the clarified supernatant was loaded onto Ni-NTA agarose beads and eluted with 50 mM HEPES, 300 mM KCl, 500 mM imidazole, 1 mM BME, pH 7.5. Eluted protein was supplemented with Ulp1 protease at the ratio of 1:50 (protease:substrate) and the digestion was conducted for 2 hours at 4 °C with mild rotation. Imidazole and salt contents were diluted with the same buffer lacking those two components, then Ulp1 and SUMO protein containing 6xHis tag were separated from DnaJB1 using Ni-NTA agarose beads. Then, DnaJB1 without any tags was further purified using ion exchange chromatography with HiTrap SP HP 5 mL column (Cytiva). Protein was eluted with a gradient of low salt (50 mM HEPES, 20 mM KCl, 1 mM BME, pH 7.0) and high salt (50 mM HEPES, 1 M KCl, 1 mM BME, pH 7.0) buffers. SDS-PAGE was used to identify fractions containing purified protein, then protein was aliquoted, flash-frozen in liquid nitrogen and stored at -80 °C until further use. Concentration of all purified proteins was measured using Pierce 660 nm Protein Assay (Thermo Scientific #22662). Purity and correct mass of all purified protein constructs were confirmed with SDS-PAGE and intact mass spectrometry. All proteins concentrations in this study refer to monomer concentration unless stated otherwise.

### Thioflavin T fluorescence aggregation assay

Prior to starting each fibrilization reaction, buffers of FL S320F tau, WT tauRD and DnaJB1 were exchanged into their fibrilization buffers using Zeba Spin Desalting Columns (Thermo Scientific #89882) and filtered with 0.22 μm centrifuge filters. This assay was conducted at 20 μM final concentration of tau protein in 100 mM potassium phosphate, 400 mM potassium citrate, 4 mM TCEP, pH 7.2 as described previously (Lövestam et al., 2025), or in 1X PBS, 10 mM DTT, pH 7.4 with equimolar (20 μM) heparin_18_ concentration (Sigma-Aldrich #H4784). DnaJB1 was added to seperate reactions to obtain a gradient of final concentrations: 20, 10, 4, 2, and 1 μM. Finally, thioflavin T (ThT; Sigma-Aldrich #596200) was filtered and added to the samples at a final concentration of 5 μM. All samples were aliquoted at 35 μL per replicate in a 384-well clear bottom plate (Corning #3544). All conditions were performed as three independent replicates at 37 °C. ThT scans were run every 10 minutes for 160 hours total, with 10 seconds of orbital shaking at 510 RPM before each read, on a Tecan Spark plate reader at 446 nm Ex (10 nm bandwidth) and 482 nm Em (10 nm bandwidth) with gain set to 65. Reactions conducted in 1X PBS with heparin were stopped after 61.8 hours due to previously observed decrease in fluorescent signal at longer incubations in these conditions. After conducted assay, endpoints were immediately flash frozen in liquid nitrogen, and stored in -80 °C until cell seeding experiments were performed. Background fluorescence signal for buffer with ThT has been subtracted from all measurements, and data plotting and estimation of t_1/2_ values were conducted in GraphPad Prism 10.2.3 using a nonlinear regression model fitting and are reported as averages with standard deviation. Cell seeding experiments for ThT assay endpoints were conducted at 100 nM final concentration (monomer concentration) for all conditions and prepared, performed and analyzed as described above for HEK293T tauRD P301S v2L FRET biosensor cells.

### Tau and α-synuclein fibrilization

Prior to starting each fibrilization reaction, buffers of all protein constructs were exchanged into their respective fibrilization buffers as indicated below and filtered with 0.22 μm centrifuge filters. Fibrilization reactions of monomeric full-length tau and tauRD with S320F point mutation were conducted as described previously (Lövestam et al., 2025) at concentrations of 60 μM and 100 μM, respectively, in 100 mM potassium phosphate, 400 mM potassium citrate, 4 mM TCEP, pH 7.2 in 1 mL total volume and at 37 °C with constant shaking at 500 RPM for 1 week. Heparin-induced fibrilization of the full-length WT tau was conducted at 8 μM protein concentration with addition of equimolar concentration of heparin_15_ (avg. 15 kDa; Amsbio #AMS.HEP001-100) in 10 mM HEPES, 100 mM NaCl, 10 mM DTT, pH 7.4, in 1 mL total volume at 37 °C for 3 days with no shaking. Fibrilization of WT α-synuclein was done as established previously (Monistrol et al., 2025) at 110 μM concentration in 50 mM sodium phosphate, 100 mM NaCl, 0.05% (w/v) NaN_3_, pH 7.3 in 1 mL total volume, at 37 °C with constant shaking at 1,000 RPM for 1 week. All fibrilization reactions were conducted in low protein binding tubes (Thermo Scientific #90410), and using Eppendorf ThermoMixer C (Eppendorf). The presence of amyloid fibrils was confirmed with positive fluorescence signal for thioflavin T and with transmission electron microscopy.

### Microscale thermophoresis (MST)

All microscale thermophoresis experiments were conducted at room temperature using a NanoTemper NT.115 (NanoTemper Technologies GmbH) with the red filter available at the Macromolecular Biophysics Resource at UTSW. Buffers of all proteins were exchanged to 1X PBS, 1 mM DTT, pH 7.4 using Zeba Spin Desalting Columns (Thermo Scientific #89882) and filtered with 0.22 μm centrifuge filters before measurements. DnaJB1 was labeled with Cyanine-5 NHS ester and final concentration of 100 nM labeled DnaJB1 and non-labeled FL tau constructs in concentration range 6 nM – 100 μM were used in all experiments. The binding measurements were prepared and conducted in the same buffer with addition of 0.05% (v/v) final concentration of Tween-20. Premium-coated capillaries (NanoTemper Technologies GmbH), and protocol of: 5 s pre-IR time, 30 s IR-on time, and 5 s of post-IR time were used. All measurements were done in three replicates, and data were analyzed using the “T-Jump” approach, with averaged replicates, and 1:1 binding model in PALMIST 1.5.8 (Scheuermann et al., 2016). A single measurement for replicate 2 of WT tau construct at concentration 24 nM has been excluded from the analysis due to irregularities in its fluorescence trace. GUSSI 2.1.6 (Brautigam, 2015) and GraphPad Prism 10.2.3 were used to visualize the results.

### Isothermal titration calorimetry (ITC)

ITC experiments were performed at 20 °C in a MicroCal PEAQ-ITC (Malvern Panalytical) calorimeter. Prior to ITC measurements, buffers of all proteins were exchanged to 10 mM HEPES, 50 mM KCl, 5 mM MgCl_2_, 2 mM TCEP, pH 7.5 using Zeba Spin Desalting Columns (Thermo Scientific #89882) and filtered with 0.22 μm centrifuge filters. To check if salt composition of the buffer impacts DnaJB1:tau_1-243_ interactions, a measurement in 50 mM sodium phosphate buffer, 2 mM TCEP, pH 6.8 (similar buffer as used in NMR) was also done as indicated in Supplementary Fig. 3A. In all experiments, the first injection had 0.5 μL volume, and was followed by twenty 1.9 μL injections, with a stirring speed of 750 RPM.

The concentrations in ITC studies for cell samples and titrants in the syringe were as follows: 20:200 μM for DnaJB1:FL WT tau, 20:200 μM for DnaJB1:WT tauRD, 20:200 μM for DnaJB1:tau_1-243_ in sodium phosphate buffer, 20:200 μM for DnaJB1:tau_1-243_ in HEPES buffer, 20:50 μM for DnaJB1:heparin_18_, 20:75 μM for DnaJB1:heparin_15_, 20:200 μM for DnaJB1:heparin_4.4_, 20:50 μM for DnaJB1:FL WT tau+heparin_18_, 20:75 μM for tau_1-243_:heparin_18_. Used heparins included: heparin_18_ (avg. 18 kDa; Sigma-Aldrich #H4784), heparin_15_ (avg. 15 kDa; Amsbio #AMS.HEP001-100), heparin_4.4_ (avg. 4351 Da; Enoxaparin sodium; Sigma-Aldrich #E0180000).

All measurements were done in two replicates, data were integrated and baseline corrected using NITPIC 2.1.0 (Keller et al., 2012). The integrated data were globally analyzed in SEDPHAT 15.2b using a model considering a single class of binding sites (Houtman et al., 2007). Binding properties were obtained from global analysis of duplicate measurements. Applied stoichiometries were deduced from the initial fits or size-exclusion chromatography experiments and were assumed as follows: 4:1 for DnaJB1:heparin_18_; 3:1 for DnaJB1:heparin_15_, DnaJB1:FL tau+heparin_18_, and tau_1-243_:heparin_18_; 2:1 DnaJB1:heparin_4.4_, and the concentrations of the syringe samples were adjusted accordingly in SEDPHAT. Correction factor for syringe species concentration was applied of 1.002 for heparin_18_. During data analysis these datapoints were excluded as their irregularity was assessed to be caused by a potential air bubble during measurement: injection 16 from replicate 1 of DnaJB1:tau_1-243_ in HEPES buffer; injections 8 and 9 from replicate 1 of tau_1-243_:heparin_18_ datasets. Thermograms and binding figures were plotted in GUSSI 2.1.6 (Brautigam, 2015).

### Size-exclusion chromatography (SEC)

DnaJB1 at 20 μM concentration in 10 mM HEPES, 50 mM KCl, 5 mM MgCl_2_, 2 mM TCEP, pH 7.5 was individually incubated with equimolar concentrations of three different species of heparin in 540 μL total volume for 30 minutes in room temperature, then centrifuged at 17,200 x g for 5 minutes, and 500 μL of obtained supernatant was injected onto pre-equilibrated Superdex 200 Increase 10/300 GL (GE) size exclusion chromatography column in the same buffer at 0.375 mL/min flow rate with measuring 215, 280, and 494 nm wavelengths. Used heparins included: heparin_18_ (avg. 18 kDa; Sigma-Aldrich #H4784), heparin_4.4_ (avg. 4351 Da; Enoxaparin sodium; Sigma-Aldrich #E0180000), and heparin-fluorescein (avg. 27 kDa; Creative PEGWorks #HP-201). Average molecular weights of heparin species were based on the information provided by the manufacturers. Standards from Gel Filtration Calibration Kit HMW (Cytiva #28403842) were prepared in the same manner as the rest of the samples and run in the same buffer.

### NMR spectroscopy

^1^H-^15^N HSQC spectra were collected at 5 °C on Agilent DD2 spectrometer operating at 800 MHz at the UTSW Biomolecular NMR Facility. Temperature of the measurement, as well as the buffer composition of 50 mM sodium phosphate, 0.01% (w/v) NaN_3_, 5% (v/v) D_2_O, pH 6.8 were selected based on the previously published data (Ukmar-Godec et al., 2020). In all experiments, ^15^N-labeled tau_1-243_ was kept constant at 25 μM with addition of DnaJB1 starting at the highest concentration (250 μM) with subsequent dilutions of the chaperone, while maintaining 25 μM concentration of tau_1-243_, with each measurement taking ∼4 hours. All NMR spectra were processed in NMRPipe (Delaglio et al., 1995) and analyzed in NMRFAM-SPARKY 1.470 (W. Lee et al., 2015). Peak assignments were transferred from the deposited data for a similar N-terminal fragment of tau protein (BMRB Entry ID 28065) (Ukmar-Godec et al., 2020). Peak intensities were determined by quantifying peak heights, and the regions of chaperone binding were identified as the regions of tau_1-243_ with signal loss greater than one standard deviation from the average intensity ratio, as established before (Irwin et al., 2021). Possible sample degradation during measurements was investigated using SDS-PAGE (Supplementary Fig. 5D). Changes in intensity ratio were plotted in GraphPad Prism 10.2.3.

### Cross-linking mass spectrometry (XL-MS)

To probe DnaJB1:tau complex formation in the presence and absence of heparin we utilized cross-linking approach with subsequent SDS-PAGE, as described before (Wydorski et al., 2023). Shortly, 3 μg of each protein were incubated together in 10 mM HEPES, 50 mM KCl, 5 mM MgCl_2_, 2 mM TCEP, pH 7.5 for 15 minutes in room temperature prior to addition of 1 mM disuccinimidyl suberate (DSS; Creative Molecules Inc. #001S). Preincubation in heparin-containing conditions was conducted with equimolar concentrations of heparin_18_ or heparin_4.4_. Cross-linking reaction was conducted for 3 minutes at 37 °C while shaking at 350 RPM and then was quenched with 50 mM ammonium bicarbonate for 10 minutes and resolved using SDS-PAGE.

To identify interaction sites in DnaJB1:tau complexes using XL-MS approach, larger scale reactions were necessary. All reactions in these experiments were conducted in 1X PBS, pH 7.4, with 15 minutes preincubation with or without equimolar concentration of heparin_18_, with cross-linking reaction conducted for three minutes at 37 °C while shaking at 350 RPM, subsequent quenching with 4X molar excess of ammonium bicarbonate, and finally resolved using SDS-PAGE. DSS was used at 1 mM final concentration, and 4-(4,6-dimethoxy-1,3,5-triazin-2-yl)-4-methyl-morpholinium chloride (DMTMM; Sigma-Aldrich #74104) was used at 43 mM final concentration. Full-length WT tau and DnaJB1 were mixed together at 20 μM concentration in 1:1 molar ratio for both DSS and DMTMM cross-linker conditions. Tau_1-243_ and DnaJB1 were mixed together at 40 μM or 50 μM concentration in 1:1 molar ratio for DSS and DMTMM, respectively. Bands relating to DnaJB1 dimer in complex with monomeric FL tau or tau_1-253_ were extracted from the gels according to the protocol established previously (Bali et al., 2023).

Mass spectrometry data were collected on Thermo Scientific Orbitrap Fusion Lumos at UTSW Proteomics Core as 5 technical replicates. Thermo RAW data files were first converted to open.mzXML format using msconvert (proteowizard.sourceforge.net), and then analyzed using locally installed xQuest 2.1.5 (Rinner et al., 2008) with FDRs estimated using xProphet (Walzthoeni et al., 2012). For DSS datasets, search parameters were defined as follows: maximum number of missed cleavages (excluding the cross-linking site) = 2, peptide length = 5–50 aa, fixed modification = carbamidomethyl-Cys (mass shift = 57.02146 Da), variable modification = oxidation of methionine (mass shift = 15.99491 Da), mass shift of the light crosslinker = 138.0680796 Da, mass shift of mono-links = 156.0786442 and 155.0964278 Da, MS1 tolerance = 10 ppm, MS2 tolerance = 0.2 Da for common ions and 0.3 Da for cross-link ions. For DMTMM datasets, search parameters were defined as follows: maximum number of missed cleavages = 2, peptide length = 5–50 residues, fixed modification = carbamidomethyl-Cys (mass shift = 57.02146 Da), variable modification = oxidation of methionine (mass shift = 15.99491 Da), mass shift of cross-linker = −18.010595 Da, no monolink mass specified, MS1 tolerance = 15 ppm, and MS2 tolerance = 0.2 Da for common ions and 0.3 Da for cross-link ions. For DMTMM datasets, results were additionally pre-filtered using an in-house shell script to identify cross-links between lysines and acidic residues. Only hits with xQuest ld-score ≥20 and FDR ≤0.05 were considered. Additionally, for each experiment, five replicate datasets were compared and only cross-link pairs that appeared in at least four datasets (DSS) or at least in three datasets (DMTMM) were interpreted. The nseen (frequency) of each residue position modified by cross-link or loop-link was summed and normalized to the total number of cross-link and loop-link modifications across all residues in each replicate and visualized using GraphPad Prism 10.2.3. The information of DnaJB1’s lysine residues that were DSS-modified (mono-links) was also extracted from the datasets to deduce about their solvent accessibility. The nseen (frequency) of each lysine residue of DnaJB1 modified by mono-link was summed and normalized to the total number of mono-link modifications across all residues in each replicate. Inter-protein cross-links were visualized using xiNET (Combe et al., 2015). Binding sites of DnaJB1 were mapped onto its dimer model made with AlphaFold2 multimer v3 (Jumper et al., 2021; Evans et al., 2021) through ColabFold (Mirdita et al., 2022; Kim et al., 2025) using PDB ID 3AGY (Suzuki et al., 2010) as a template and then visualized in ChimeraX 1.9. Potential heparin-binding sites on DnaJB1 dimer were visualized using CTD-DD-containing structure under PDB ID 3AGY. For testing whether heparin presence increases the number of found inter-protein cross-links in our datasets, unpaired t tests were used to assess statistical significances relative to no heparin controls using GraphPad Prism 10.2.3.

## Data and code availability

All cell-based aggregation, ThT, ITC, NMR, MST, SEC, and XL-MS data, as well as uncropped western blot membranes and SDS-PAGE gels are available as Source Data. Source Data, plasmid maps of constructs used in this study, AlphaFold model of full-length DnaJB1 dimer and raw data associated with this work are available on Zenodo under accession number 17228160. Raw MS data used for the XL-MS analysis are available in the MassIVE database under the accession number [MSV000099233](https://doi.org/10.25345/C5610W516).

NMR signal assignments for tau_1-243_ used in this study were based on BMRB Entry ID 28065. Model of full-length DnaJB1 dimer was made with AlphaFold2 multimer v3 based on PDB ID 3AGY. PALMIST, NITPIC, and GUSSI software used for MST and ITC data analysis and visualization are available on the UTSW Macromolecular Biophysics Resource website [https://www.utsouthwestern.edu/research/core-facilities/mbr/software/]. ITC data were also analyzed using SEDPHAT [https://sedfitsedphat.github.io/sedphat/download.htm]. NMR spectra were processed using NMRPipe [https://www.ibbr.umd.edu/nmrpipe/install.html] and analyzed in NMRFAM-SPARKY [http://pine.nmrfam.wisc.edu/download_packages.html]. XL-MS data were analyzed using xQuest [https://gitlab.ethz.ch/leitner_lab/xquest_xprophet]. Cross-links were visualized using xiNET [http://crosslinkviewer.org/]. Full-length DnaJB1 dimer was made using ColabFold [https://protocol.colabfold.com].

## Acknowledgements

We thank Shih-Chia (Scott) Tso and Chad Brautigam in the UTSW Macromolecular Biophysics Resource for help with MST and ITC training and data analysis. We appreciate the help of the Proteomics Core Facility at UTSW with mass spectrometry data collection. NMR data was acquired at the Biomolecular NMR Facility at UTSW. We would like to also thank Dominika Borek for the suggestion to include low-molecular-weight heparin in our experiments. This research was supported by NIH-NIA 1RF1AG078888-01 and NIH-NINDS 1R01AG083876 grants to LAJ and an award from The Carl B. & Florence E. King Foundation to PMW. This paper was typeset with the template by @Chrelli: www.github.com/chrelli/bio-Rxiv-word-template.

## Author contributions

PMW and LAJ initiated this project. PMW designed, conducted, and analyzed all experiments. PM contributed to protein purification and sample preparation for cross-linking mass spectrometry. SB contributed to NMR data collection and analysis. JVA contributed to N-terminal tau truncation experiment design, obtaining plasmids, and data analysis. PMW and LAJ acquired funding and supervised this project. PMW and LAJ prepared the original draft, and all authors contributed to the review and editing of the final version of this manuscript.

## Competing interest statement

JVA is a co-founder of Handshake Bio. Handshake Bio did not directly fund or influence the design, execution, or interpretation of the experiments presented in this manuscript. The remaining authors declare no competing interests.

## Supplementary Material for

**Supplementary Figure 1.**
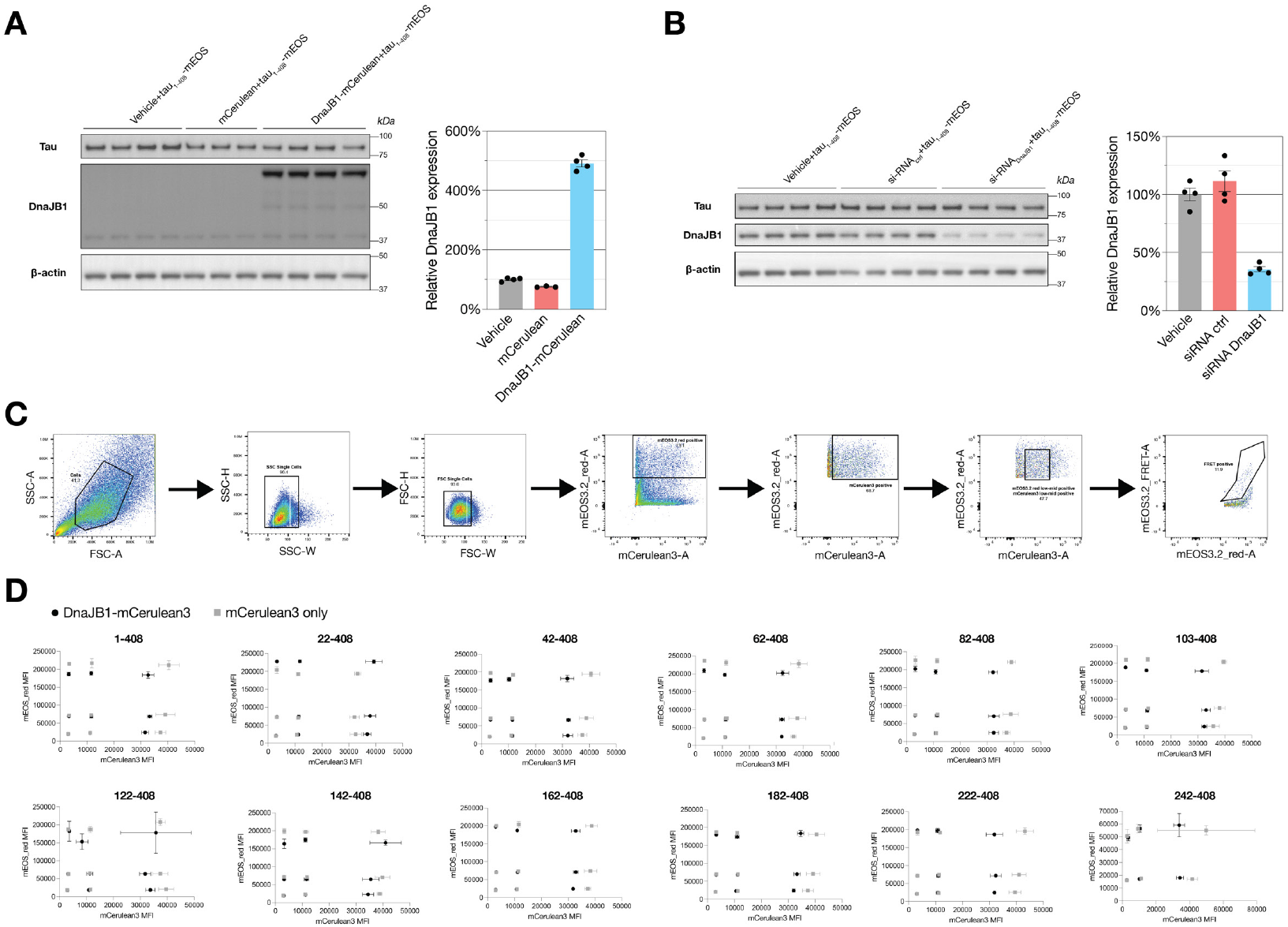
Cell biosensor model allows to investigate the role of DnaJB1 chaperone in tau aggregation. **(A)** Western blot analysis of DnaJB1-mCerulean3 overexpression with subsequent overexpression of the longest tau construct tested (1-408) at the time point of tau seed addition in HEK293T cells, 4 days after DnaJB1-mCerulean3 or mCerulean3-only transfection. On the right is the normalized quantification of the DnaJB1 expression levels. Data are shown as averages with error bars representing SEM across 4 (for Vehicle and DnaJB1-mCerulean3 overexpression conditions) or 3 (for mCerulean3 overexpression control condition) independent transfections. **(B)** Western blot analysis of DnaJB1 knockdown with siRNA duplex oligo approach with subsequent overexpression of the longest tau construct tested (1-408) at the time point of tau seed addition in HEK293T cells, 4 days after siRNA transfection. On the right is the normalized quantification of the DnaJB1 expression levels. Data are shown as averages with error bars representing SEM across 4 independent transfections. **(C)** Gating strategy used in HEK293T cells experiments to investigate tau aggregation of different N-terminal truncations of tau protein under overexpression or knockdown of DnaJB1 chaperone. Populations representing live, single cells were isolated with subsequent separation of the cells positive for mEOS3.2 red fluorescent signal. Then, in case of the experimental setup containing DnaJB1 overexpression with mCerulean3 fluorescent tag, additional gate was used to exclude the highest mCerulean3-expressing population. The highest mEOS3.2-expressing populations were excluded due to the discrepancy of mEOS3.2 red vs mCerulean3 fluorescence for the highest expressing populations as shown in panel (D). **(D)** Median fluorescent intensity (MFI) of red fluorescence of mEOS3.2 and fluorescence of mCerulean3 for all tested constructs of N-terminus truncation of tau protein tagged with mEOS3.2 in the context of DnaJB1 overexpression with mCerulean3 tag. Each population was divided into 3 distinct gates representing their low, medium, or high expression in either of the channels. Black symbols represent DnaJB1-mCerulean3 overexpression, and gray symbols represent mCerulean3 only control. Error bars represent 3 biological replicates.

**Supplementary Figure 2.**
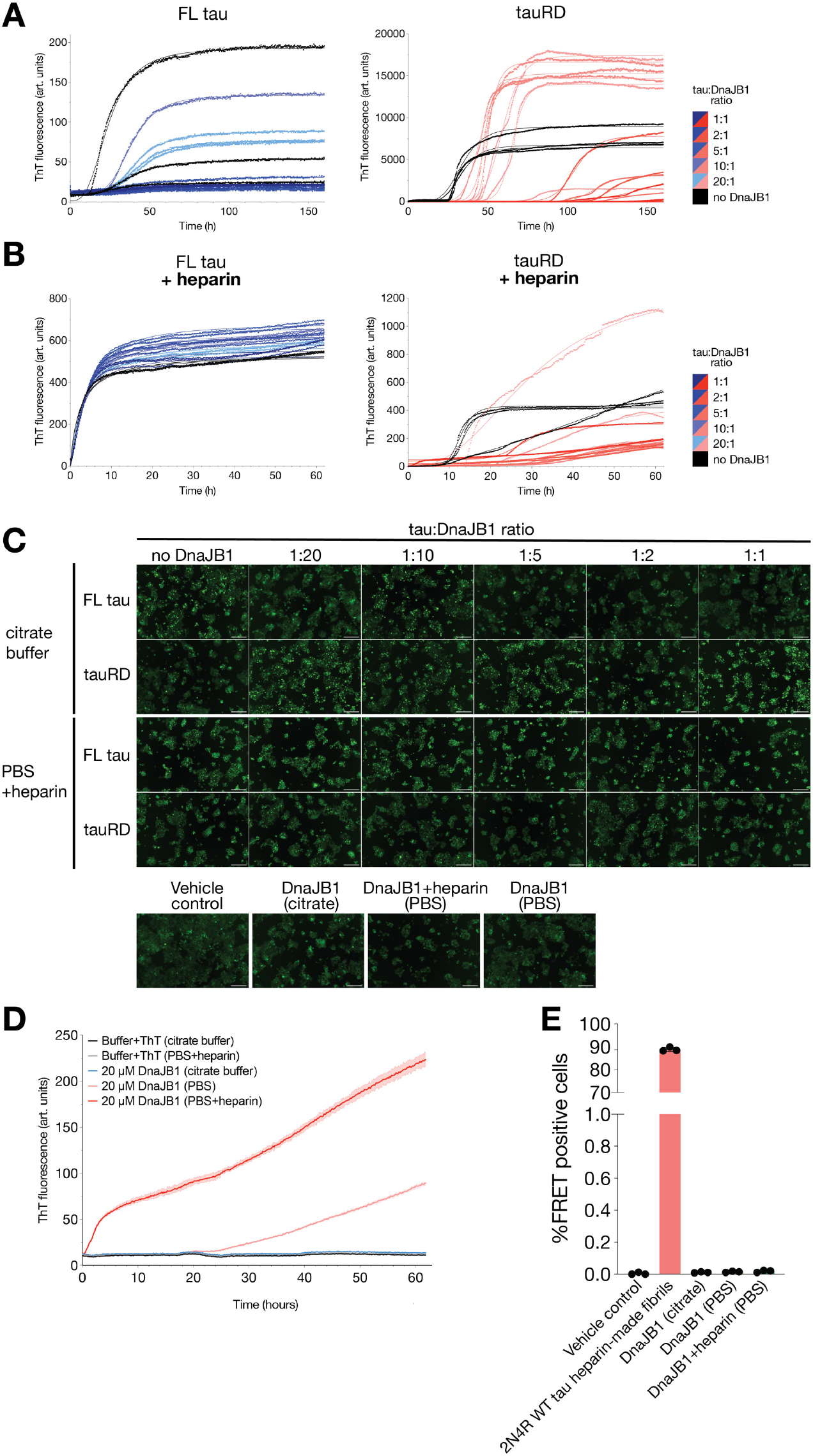
Concentration dependent effect of DnaJB1 on tau aggregation investigated with ThT fluorescence aggregation, and cell seeding assays. **(A)** Nonlinear regression fitting of thioflavin T (ThT) fluorescence signals of FL S320F tau (left, blue) and WT tauRD (right, red) incubated in absence of DnaJB1 (black) or with different concentrations of DnaJB1 (color gradients) in 100 mM potassium phosphate, 400 mM potassium citrate, 4 mM TCEP, pH 7.2. Each condition with different chaperone concentration was done in triplicate, symbols represent ThT fluorescence measurements and solid line represent nonlinear regression fitting. **(B)** Nonlinear regression fitting of thioflavin T (ThT) fluorescence signals of FL S320F tau (left, blue) and WT tauRD (right, red) incubated in absence of DnaJB1 (black) or with different concentrations of DnaJB1 (color gradients) in 1X PBS, 10 mM DTT, pH 7.4 with equimolar concentration of heparin. Each condition with different chaperone concentration was performed in triplicate, symbols represent ThT fluorescence measurements and solid line represent nonlinear regression fitting. **(C)** Representative fluorescent microscope images of cell seeding experiments of ThT aggregation assay endpoints of FL S320F tau and tauRD aggregated with different concentrations of DnaJB1 in buffers with or without heparin (as shown in Figure 2EI). Tau biosensors were seeded with 100 nM final concentration of the ThT assay endpoints and after 48 hours the development of tau inclusions was investigated. Scale bars represent 200 μm. **(D**) ThT fluorescence signals of 20 μM DnaJB1 incubated in different buffers: potassium citrate buffer (blue), 1X PBS (light red), 1X PBS with equimolar concentration of heparin_18_ (dark red). Each buffer condition was done in three replicates, solid lines represent ThT fluorescence measurements and error lines represent the ranges of replicates. **(E)** Quantification of tau inclusion formation when tau biosensor cells were seeded with 100 nM of the ThT endpoint of DnaJB1 incubated in different buffers. Despite ThT-positive signal of DnaJB1 incubated in PBS and in PBS with heparin, these reactions did not lead to emergence of seeding-competent species. Data are shown as averages with error bars representing SD across 3 independent transfections.

**Supplementary Figure 3.**
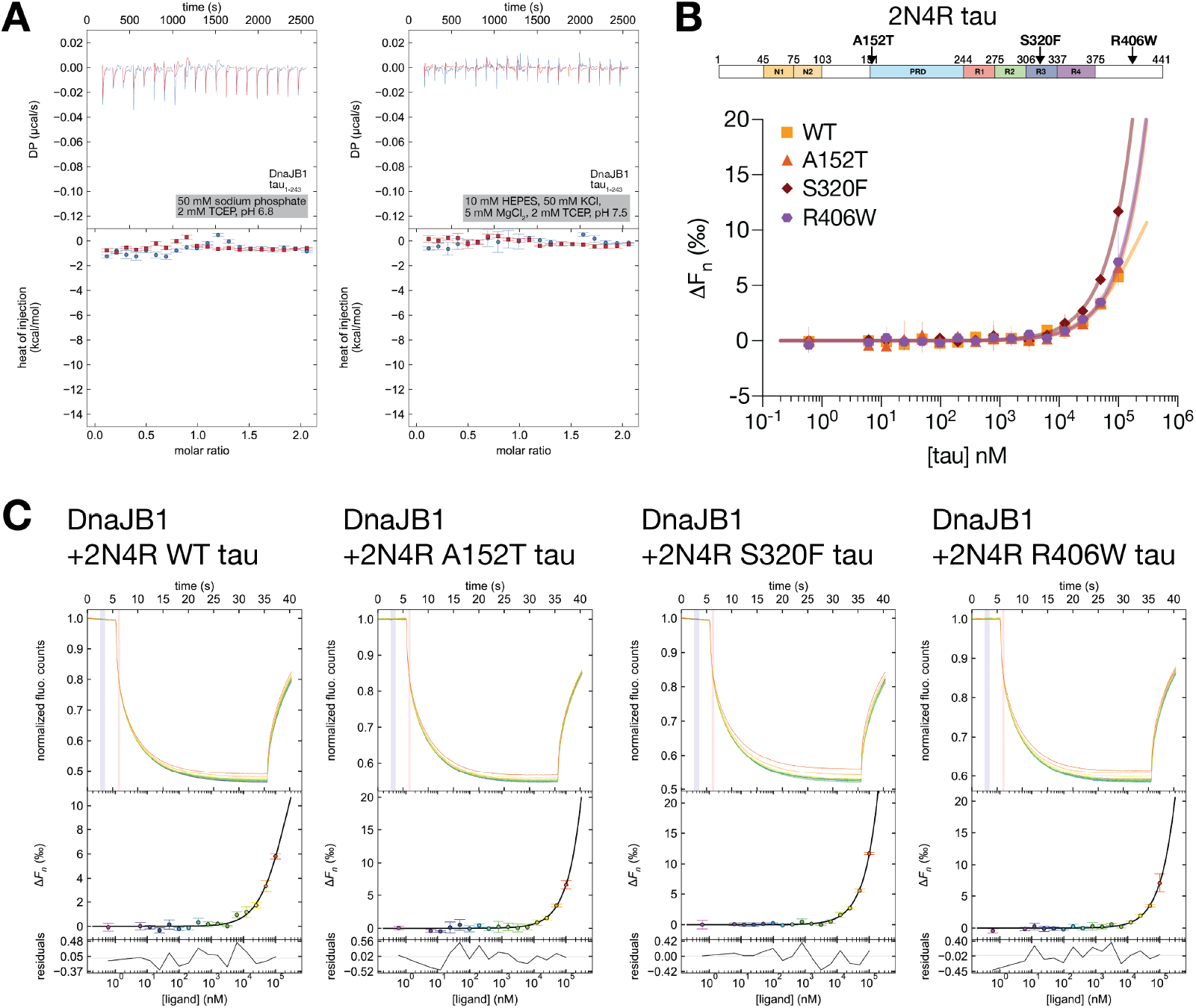
DnaJB1 does not bind monomeric forms of 2N4R tau in the absence of heparin. **(A)** ITC titration of DnaJB1 binding to tau_1-243_ in two different buffers: 50 mM sodium phosphate buffer, 2 mM TCEP, pH 6.8 or 10 mM HEPES, 50 mM KCl, 5 mM MgCl_2_, 2 mM TCEP, pH 7.5 showing that changes in salt composition and concentration in the used buffer do not enhance DnaJB1:tau binding. The top panels show reconstructed thermograms and the bottom panels show isotherms with no detectable binding. **(B)** MST titrations of DnaJB1 with 2N4R tau constructs WT (orange squares), or disease-associated point mutants: A152T (red triangles), S320F (dark red rhombi), R406W (purple hexagons). Fluorescence change (ΔF) is plotted as a function of 2N4R tau concentration. Data are shown as triplicates and are plotted as the average with the range of individual replicates. Data were fitted to the 1:1 binding model using PALMIST. **(C**) Cumulative raw thermophoresis data for three independent titration experiments summarized in the panel (B). The curves represent normalized fluorescence counts colored by concentration of 2N4R tau from blue (low) to red (high) and plotted as a function of time. Fluorescence change (ΔF) is plotted as a function of substrate concentration colored as above with the black line representing the fit to the 1:1 binding model using PALMIST. Data are shown as averages with SD across three independent experiments. The variance between the average ΔF value and the fit is plotted below. The residuals are shown as average deviation from the fit across three experiments.

**Supplementary Figure 4.**
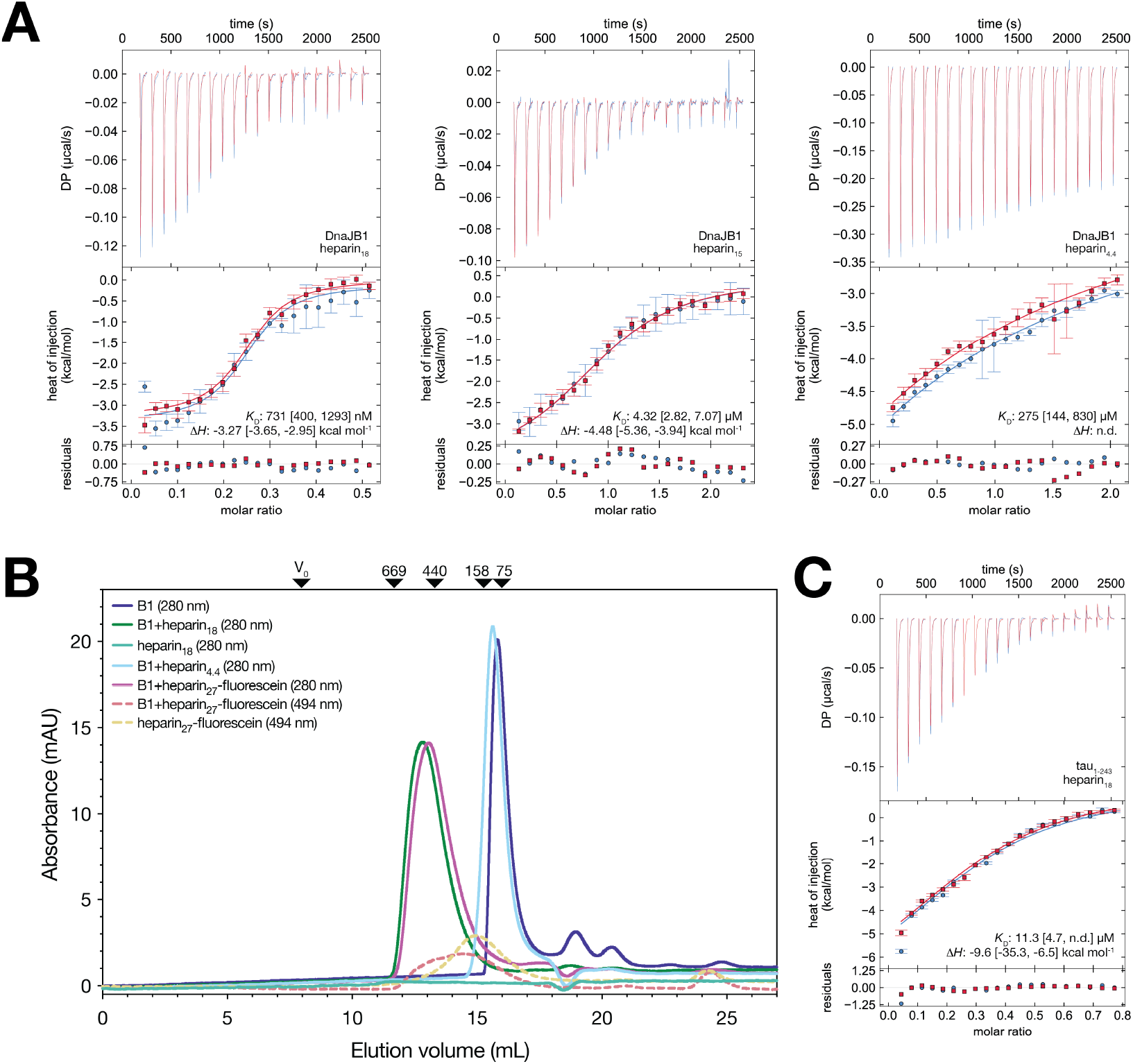
Multivalency dependence of heparin binding. **(A)** ITC titrations of DnaJB1 binding to different sizes of heparin: heparin with average mass of ∼18 kDa (left), ∼15 kDa (middle), and ∼4.4 kDa (right). The top panels show reconstructed thermograms from NITPIC, the middle panels show binding isotherms and individual fits, and the bottom panels show the fitting residuals. Error bars on reconstructed thermograms represent estimated peak integration errors from NITPIC. Square brackets represent a 68.3% confidence interval determined by error surface projection in global analysis of duplicate experiments in SEDPHAT. n.d. not determined. **(B)** Size-exclusion chromatography (SEC) of DnaJB1 incubated with different sizes of heparin with measuring absorbance in 280 nm (solid lines) or in 494 nm (dotted lines): DnaJB1 alone (dark blue), DnaJB1 with heparin avg. 18 kDa (dark green), heparin avg. 18 kDa alone (light green), DnaJB1 with heparin avg. 4.4 kDa (light blue), DnaJB1 with heparin-fluorescein avg. 27 kDa (violet solid line represents absorbance signal in 280 nm, orange dotted line represents absorbance signal in 494 nm), heparin-fluorescein avg. 27 kDa alone (yellow dotted line represents absorbance signal in 494 nm). Black triangles represent the maximum absorbance signal for calibration standards and numbers refer to their molecular weights in X·10_3_, as indicated by the producer. **(C)** ITC titrations of tau_1-243_ binding to heparin species with average mass of ∼18 kDa. Top, middle, and bottom panels as in (A).

**Supplementary Figure 5.**
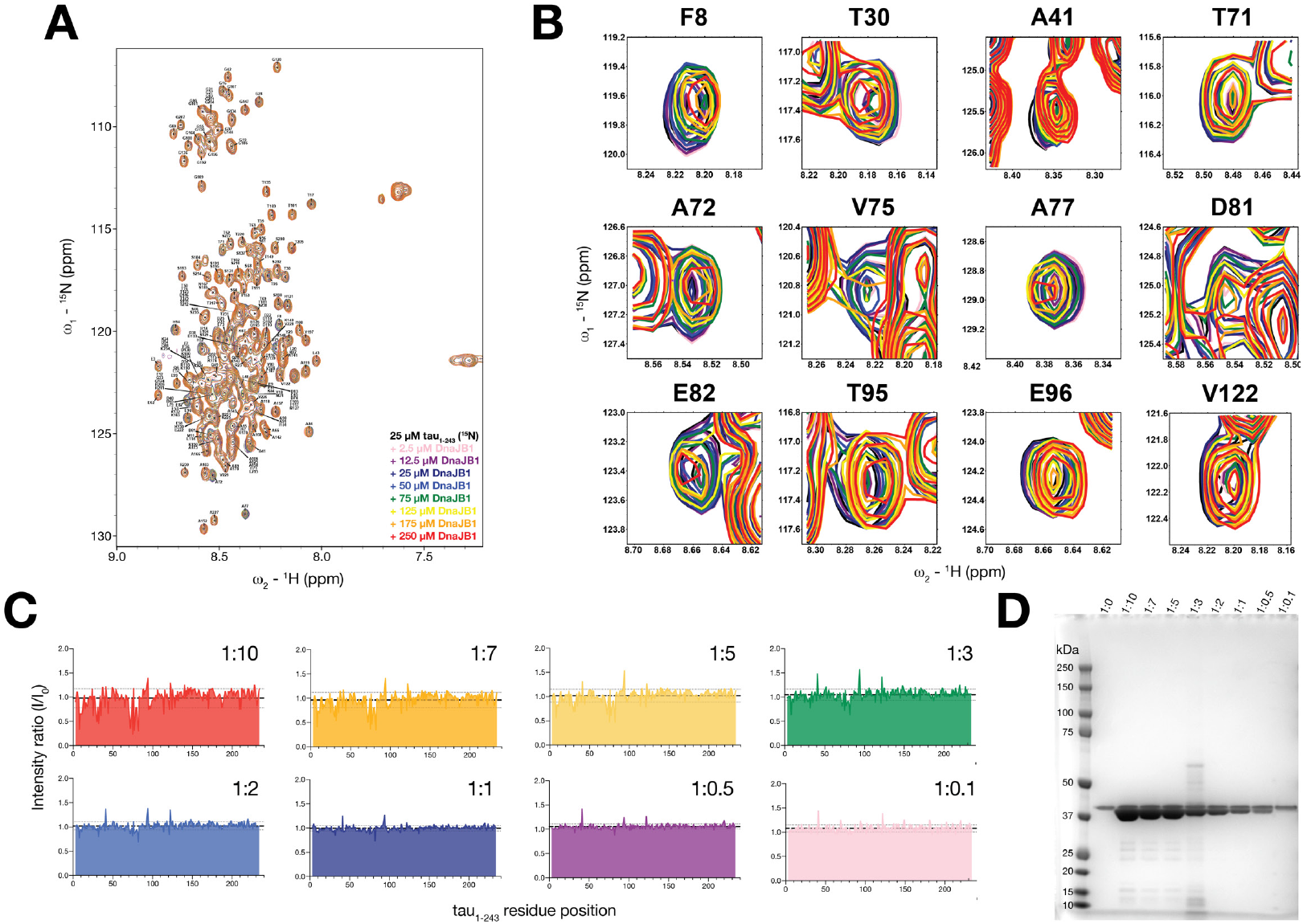
DnaJB1 chaperone binds to three distinct sites on the N-terminus of tau. **(A)** Superposition of _1_H-_15_N HSQC spectra of tau_1-243_ in the absence (black) and different concentrations of DnaJB1 as indicated by the color code (DnaJB1 concentrations gradually decreased from 250 to 2.5 μM). Peak assignment was taken from the previously deposited BMRB Entry 28065. **(B)** Expansions showing the changes observed at increasing DnaJB1 concentrations in selected labeled cross-peaks from tau_1-243_. The color code is the same as in (A). **(C)** Summary of the peak intensity ratio I/I_0_ of tau_1-243_ (I_0_) and tau_1-243_ with DnaJB1 (I) at different tau_1-243_:DnaJB1 molar ratios. Thin dotted lines represent the range of single standard deviation from the average intensity ratio of tau_1-243_ in the absence of DnaJB1 (bold dashed line). The color code is the same as in (A). **(D)** SDS-PAGE gel of samples used in _1_H-_15_N HSQC NMR experiments after spectra collection to investigate possible sample degradation/aggregation.

**Supplementary Figure 6.**
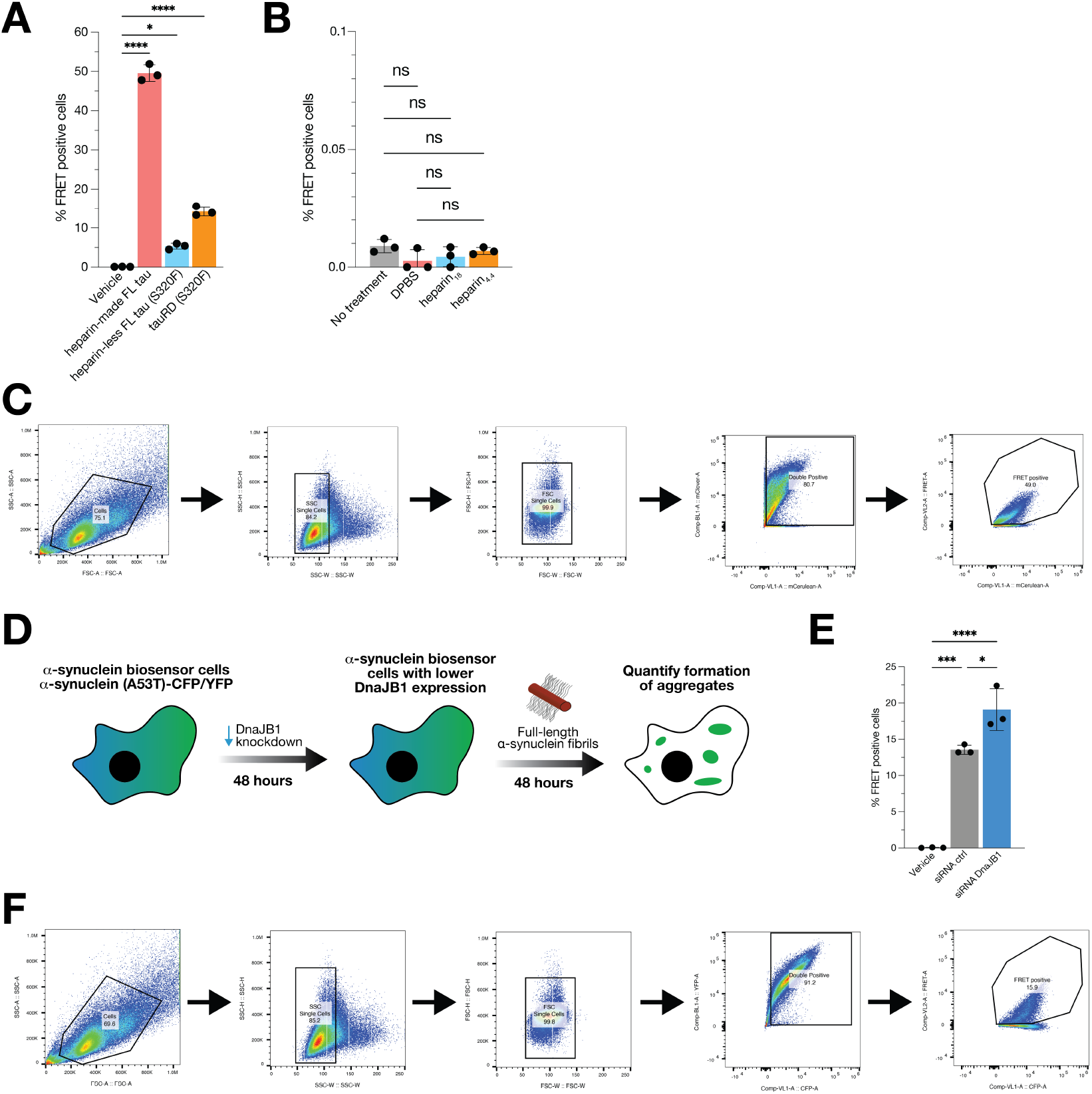
DnaJB1 knockdown increases α-synuclein aggregation in cell biosensors. **(A)** Summary of seeding efficiencies of all recombinant fibrils at 100 nM final concentration used in this manuscript with tau fibrils tested in HEK293T tauRD P301S v2L FRET biosensor cell line. Data are shown as averages with error bars representing SD across 3 independent transfections. (*) p < 0.05, (****) p < 0.0001. **(B)** Treatment with different species of heparin does not induce tau aggregation in tauRD cell biosensors. Data are shown as averages with error bars representing SD across 3 independent transfections. ns=not significant. **(C)** Gating strategy used in tauRD biosensor cells stably expressing tau_246-378_ (P301S) with mCerulean3 or mClover3 fluorescent tag. Populations representing live, single cells were isolated with subsequent separation of the cells double positive for mCerulean3 and mClover3 fluorescent signal. Then, a FRET signal was measured as the percentage of mClover3-mCerulean3 FRET positive cells in the parent population (double positive). **(D)** Schematic of the experimental setup to investigate DnaJB1 knockdown in biosensor cells stably expressing α-synuclein (A53T) with either CFP or YFP fluorescent tag. α-synuclein aggregation was measured as a FRET signal via flow cytometry 48 hours after seeding with recombinant α-synuclein fibrils. **(E)** Lipofectamine-mediated α-synuclein seeding in α-synuclein biosensor cells in the context of DnaJB1 knockdown. Data are shown as averages with error bars representing SD across 3 independent transfections. (*) p < 0.05, (***) p < 0.001, (****) p < 0.0001. **(F)** Gating strategy used in α-synuclein cell biosensors cells stably expressing α-synuclein (A53T) with CFP or YFP fluorescent tag. Populations representing live, single cells were isolated with subsequent separation of the cells double positive for CFP and YFP fluorescent signal. Then, a FRET signal was measured as the percentage of CFP-YFP FRET positive cells in the parent population (double positive).

**Supplementary Figure 7.**
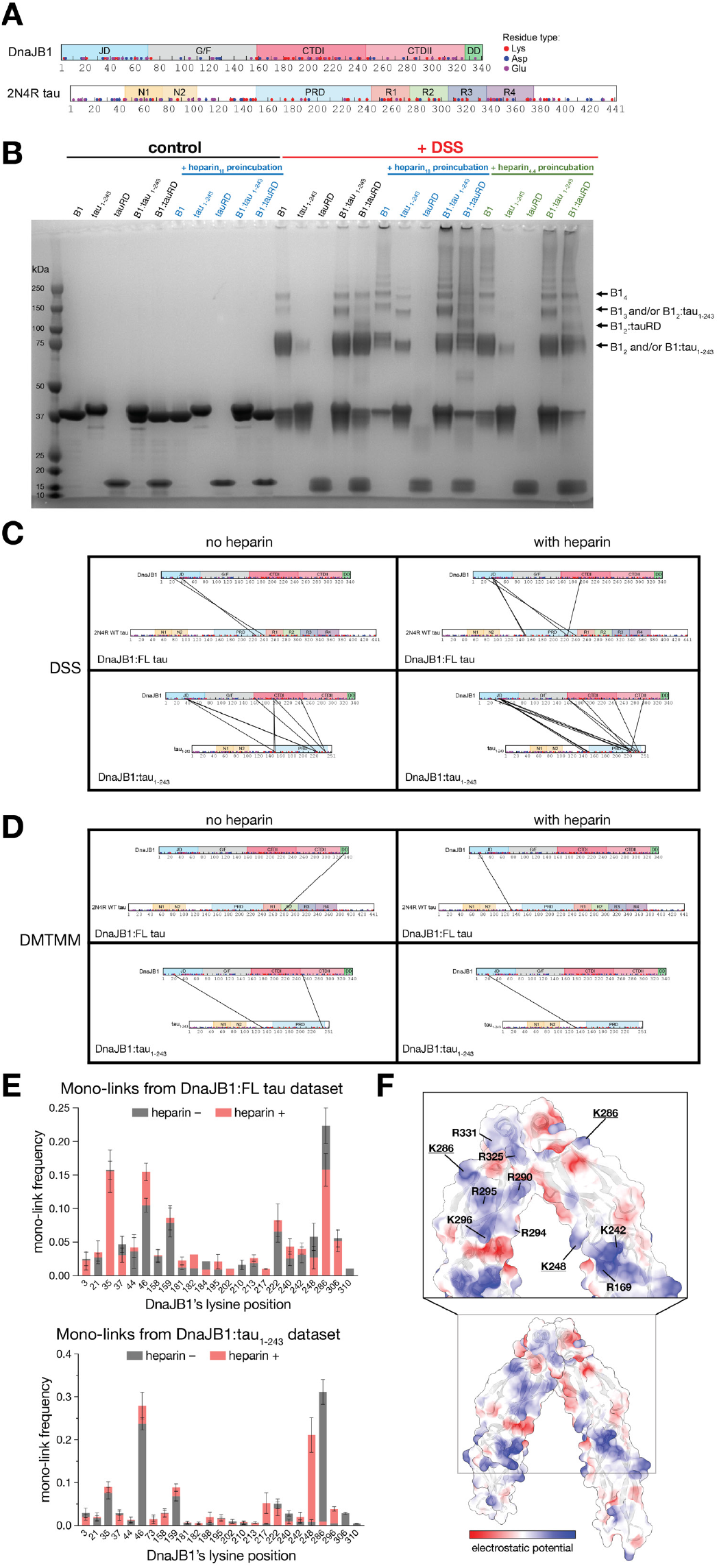
Cross-linking mass spectrometry (XL-MS) investigation of DnaJB1:tau heterocomplexes. **(A)** Domain architectures of human DnaJB1 and 2N4R isoform of tau protein. Domains of DnaJB1 labeled and colored as in Fig. 5C, and domains of 2N4R tau labeled and colored as in Fig. 1A. With red, blue, and violet spheres are indicated positions of lysine, aspartic acid, and glutamic acid residues, respectively, as these are the residues accessible to be modified by the cross-linkers used in this study. **(B)** Cross-linked samples reveal the formation of covalent heterocomplex bands (red, DSS cross-linker) compared to untreated reactions (black, control) by SDS-PAGE. Preincubation of protein samples with heparin with average mass of ∼18 kDa seems to enhance heterocomplex formation (blue), but less so if preincubation was led in the presence of smaller ∼4.4 kDa heparin (green). **(C)** Schematic showing identified inter-protein cross-links in heterocomplexes of DnaJB1 with FL 2N4R tau (top) or tau_1-243_ (bottom) using DSS cross-linker (Lys-Lys contacts). XL-MS experiments were conducted in the absence (left) or in the presence of heparin (right). Only inter-protein cross-links that appeared among 4 or more of 5 total technical replicates were visualized. **(D)** Schematic showing identified inter-protein cross-links in heterocomplexes of DnaJB1 with FL 2N4R tau (top) or tau_1-243_ (bottom) using DMTMM cross-linker (Lys-Asp or Lys-Glu contacts). XL-MS experiments were conducted in the absence (left) or in the presence of heparin (right). Only inter-protein cross-links that appeared among 3 or more of 5 total technical replicates were visualized. **(E)** Quantification of identified mono-links of DnaJB1 from DSS XL-MS datasets with FL tau (top) or tau_1-243_ (bottom) as clients. Data are shown as the nseen (frequency) of each lysine residue of DnaJB1 modified by mono-link that was normalized to the total number of mono-link modifications across all residues in each replicate. Data are shown as the averages of the normalized frequencies with SD across 5 technical replicates. **(F)** Basic patches of the β-sandwich C-terminal domains of DnaJB1 dimer that present as the potential heparin-binding sites. Secondary structure of DnaJB1 dimer (with only CTDI/II–DD present in the deposited model) from PDB ID 3AGY (Suzuki et al., 2010) in cartoon representation and the electrostatic potential is represented as surface in red-white-blue gradient.

## Notes

https://doi.org/10.5281/zenodo.17228160

